# Fecal *Dysosmobacter* spp. concentration is linked to plasma lipidome in insulin-resistant individuals with overweight, obesity and metabolic syndrome

**DOI:** 10.1101/2025.03.19.644117

**Authors:** Camille Petitfils, Clara Depommier, Nathalie M. Delzenne, Amandine Everard, Matthias Van Hul, Patrice D. Cani

## Abstract

**Background:** Obesity is reaching epidemic proportions worldwide. This excessive increase of adipose tissue is a risk factor for the development of multiple diseases and premature death. Amongst associated diseases, metabolic syndrome is one of the main comorbidities of obesity. In this context, the gut microbiota has been recognized as both shaping and responding to host energy metabolism. Recently metabolomics has emerged as a powerful tool to capture a snapshot of the metabolites present in a specific tissue, offering new insights into host-microbiota interactions. Integrating metabolomics with gut microbiota studies could help us better understand how specific species impact on host metabolomic profile. *Dysosmobacter welbionis* has been identified as a promising next generation beneficial bacteria with potential effects on fat mass and glucose metabolism in mice, and fecal *Dysosmobacter spp* concentration was inversely correlated to body mass index fasting glucose and plasmatic HbA1c in humans.

**Methods:** Concentration of *Dysosmobacter spp* was quantified by qPCR in the stools of insulin resistant overweight/obese participants with a metabolic syndrome and plasma metabolites were analyzed using untargeted metabolomics. Correlations between *Dysosmobacter spp* fecal abundance and the 1169 identified plasma metabolites were uncovered using Spearman correlations followed by a false discovery rate correction.

**Results:** Interestingly, among the detected metabolites, *Dysosmobacter spp* was exclusively associated with lipid molecules, primarily structural lipids involved in membrane formation. This finding aligns with previous *in vivo* studies highlighting lipid profile alterations in multiple tissues of mice treated with this bacterium.

**Conclusion:** These results suggest that *Dysosmobacter spp* plays a specific role in host lipid metabolism. Further studies are needed to elucidate the underlying mechanisms and assess its potential therapeutic applications.

## Introduction

In its most recent report, the World Health Organization established that 1 in 8 people in the world now lives with obesity. This number rises to nearly three billion people when overweight individuals are included. While obesity is not always a direct predictor of metabolic health (1), central obesity (or abdominal) obesity is a key diagnostic criterion for metabolic syndrome.

Metabolic syndrome is defined by meeting at least three of the five following criteria: central obesity, hypertension, hypertriglyceridemia, low levels of HDL cholesterol and elevated glycemia (2). Consequently, living with a metabolic syndrome significantly increases the risk of developing type 2 diabetes (T2D) and cardiovascular diseases (2). Obesity and its comorbidities stem from complex and multifactorial causes, prompting significant efforts to understand how they interact and how to tackle this growing epidemic. Over the last decades, the gut microbiota has emerged as a key player in host pathophysiology, influencing the development of metabolic diseases. Recently, several studies have demonstrated the therapeutic potential of specific gut bacteria. For instance, in men with metabolic syndrome, a four-week treatment with *Anaerobutyricum soehngenii* improved glucose metabolism (3). Similarly, in the Microbes4U^®^ cohort, administration of pasteurized *Akkermansia muciniphila* helped mitigate the progression of cardiometabolic disorders associated with overweight and obesity (4).

In 2020, a novel bacterial species, *Dysosmobacter welbionis* J115^T^, was isolated. Two years later, this bacterium was associated with beneficial effects on weight and glucose tolerance in a mouse model of obesity (5, 6). In human, *Dysosmobacter spp* fecal concentrations was found to be negatively correlated with body mass index (BMI), fasting glucose and plasmatic HbA1c (5) in a cohort of overweight or obese people with a metabolic syndrome and insulin resistance. In a randomized trial (Food4Gut), *Dysosmobacter spp* fecal concentration was unaffected by prebiotic (inulin) supplementation but obese/diabetic subjects who responded best to the prebiotic intervention had higher baseline *D. welbionis* levels than non-responders, and *D. welbionis* was negatively correlated with fasting glycemia. Interestingly, *D. welbionis* was significantly more abundant in the feces of metformin treated patients (7), although *in vitro* experiment did not reveal a direct utilization of metformin by this bacterium.

Metabolomics has become a valuable tool for deciphering metabolic disturbances in both health and disease (8, 9). By capturing a real-time snapshot of the metabolites within a tissue, it provides critical insights into underlying pathological processes and treatment responses, ultimately contributing to the advancement of personalized medicine. Given the intricate relationship between gut microbiota and host health, more and more studies are exploring how the gut microbiota composition correlates with host metabolomic profiles (10, 11). Blood metabolites, in particular, hold significant potential as they can be assessed through minimally invasive sampling. Although the origin of many metabolites remains unclear, studies consistently highlight the gut microbiota’s influence on blood metabolomic composition (12–17).

As *Dysosmobacter spp* has been associated with an improved phenotype in people living with overweight/obesity and metabolic syndrome, this study aims to explore its association with plasma metabolites in this population. Identifying a specific metabolomic profile associated with *Dysosmobacter spp* abundance in overweight humans could enhance our understanding of its role in host metabolic health.

## Methods

### Data source and study participants

The metabolic parameters, metabolomic profile and fecal concentration of *Dysosmobacter spp* were obtained from non-diabetic, insulin-resistant and overweight or obese individuals previously recruited in the Microbes4U cohort (4, 5, 18, 19). Briefly, participants were recruited at the Cliniques Universitaires Saint-Luc located in Brussels, Belgium, between 2015 and 2018. 52 overweight or obese subjects (body mass index > 25 kg/m^2^) newly diagnosed with a metabolic syndrome and with a prediabetes state as well as an insulin sensitivity < 75% were included. Metabolic syndrome was diagnosed according to the National Cholesterol Education Program Adult Treatment Panel III definition, that is, at least three of the five following criteria: fasting glycemia > 100 mg/dL; blood pressure ≥ 130/85 mmHg or antihypertensive treatment; fasting triglyceridemia ≥ 150 mg/dL; high-density lipoprotein (HDL) cholesterol < 40 mg/dL for men, <50 mg/dL for women; and/or waist circumference > 102 cm for men, >88 cm for women.

Insulin sensitivity and resistance were both analyzed in Depommier et al 2019 (4) using HOMA. Three blood samples, at 5 min intervals, were taken for each individual. Insulinemia and glycemia were determined for each sample and the mean values were then entered in the HOMA2 calculator (v.2.3.3, available from http://www.dtu.ox.ac.uk/homacalculator/) to estimate insulin sensitivity (%) and insulin resistance (20, 21).

Anthropometric measurements and plasma metabolomics were performed in Depommier et al 2019 (4). Bodyweight (BW) was measured in kg, BMI in kg.m^−2^, and waist and hip circumferences in cm using a flexible tape. After an overnight fasting of 8 hours minimum, blood samples were collected in different tubes: sodium fluoride-coated tubes for fasting glycemia and insulinemia; lipopolysaccharide (LPS)-free heparin sulfate coated tubes for LPS measurement (BD Vacutainer, NH sodium heparin, 368480) measured by Endosafe-MCS (Charles River Laboratories, Lyon, France) in Depommier et al 2019 (4); lithium-heparin-coated tubes for enzymatic activities. Fasting glycemia, insulinemia, HbA1c (%), total cholesterol, LDL cholesterol (calculated), HDL cholesterol, triglycerides (TG), gamma-glutamyl transferase (GGT), alanine transaminase (ALT), aspartate transaminase (AST), lactate dehydrogenase (LDH), creatine kinase (CK), creatinine, glomerular filtration rate (GFR), urea, C-reactive protein (hsCRP), alkaline phosphatase (AlkP), and, white blood cell count (WBC) were measured directly at the hospital in the same paper (4). Plasma non-esterified fatty acids (NEFA) were measured in Depommier et al 2021(18), using kits coupling an enzymatic reaction with spectrophotometric detection of the reaction end products (Diasys Diagnostic and Systems, Holzheim, Germany).

### Metabolomic analysis

Plasma was isolated by centrifugation at 4200g for 10min at 4°C and stored at −80°C. 100µl was aliquoted for metabolomics analyses by Metabolon Inc (North Carolina, USA). Untargeted metabolomics analyses were performed as described in Babu et al 2019 (22). It comprised ultra-performance liquid chromatography/mass spectrometry with a heated electrospray ionization source and mass analyzer. Following proper handling, samples were first prepared using the automated MicroLab STAR® system from Hamilton Company. The resulting extract was divided into several fractions, analysed by four ultra-high-performance liquid chromatography-tandem mass spectrometry according to the Metabolon pipeline. Biochemical identification of metabolites contained in one sample was then performed by comparison to a reference library of purified standards consisting of more than 33000 metabolites. The comparison was based on retention time/index, mass-to-charge ratio (m/z), and chromatographic data (MS/MS spectral data) using software developed at Metabolon. Further details regarding quality controls, data extraction, curation, quantification, and bioinformatics were previously described (23). Compounds that have not been confirmed based on a standard, but that are identified by Metabolon Inc with confidence are labelled with an asterisk (*). Structural isomers of another compound in Metabolon Inc spectral library are labelled with the isomer position number in brackets ([#]).

### Plasma biomarkers

Plasmatic monocyte chemoattractant protein 1 (MCP-1) was assessed in Depommier et al 2019 (4) and plasmatic leptin in Depommier et al 2021 (19). Plasmatic interferon gamma-induced protein 10 (IP-10), soluble CD40 ligand (sCD40L), eotaxin, growth differentiation factor 15 (GDF-15), soluble vascular cell adhesion molecule-1 (svCAM-1), lipocalin-2, soluble intercellular adhesion molecule-1 (siCAM-1), ADAMts13, D-Dimer, myoglobulin, glucagon, gastric inhibitory polypeptide (GIP) and macrophage-derived chemokine (MDC) were measured using the same MILIPLEX MAP Human Metabolic Hormone Magnetic Bead Panel technology and measured using Luminex technology (BioPlex; Bio-Rad Laboratories, Hercules, CA, USA) according to the manufacturer’s instructions.

Active plasma glucagon-like-peptide-1 (GLP-1) was measured in Depommier et al 2019 (4) by sandwich ELISA (Merck Millipore). Plasmatic GLP-2, peptide YY (PYY), intestinal fatty-acid binding protein (I-FABP), and interleukin 6 (IL-6) were assessed using the same method, following the manufacturer’s instructions. Dipeptidyl peptidase 4 (DDP4) activity was measured previously (4), through the quantification of p-nitroanilide (pNA) production from glycine-proline-pNA (Sigma-Aldrich).

### Fecal Dysosmobacter quantification

The quantity of *Dysosmobacter spp* per gram of feces for each patient upon recruitment has been measured in Le Roy et al 2022 (5). Concisely, genomic DNA was extracted from human stools using the QIAamp DNA Stool Mini Kit (Qiagen, Germany), including a bead-beating step. Quantified standard DNA for *Dysosmobacter spp* qPCR was obtained by extracting genomic DNA with the same DNA extraction protocol than for the stools from a *Dysosmobacter welbionis* J115^T^ culture in exponential growing phase of known concentration in colony forming units (CFU) determined by plating. DNA concentration was determined, and purity (A260/A280) was checked using a NanoDrop2000 (Thermo Fisher Scientific, USA). Samples were diluted to an end concentration of 10 and 0.1 ng/µl in TE buffer pH 8. Total bacteria qPCR was performed on the 0.1 ng/µl dilution and the *Dysosmobacter* spp qPCR was performed on the 10 ng/µl dilution. Real-Biosystems, Den Ijssel, The Netherlands) using Mesa Fast qPCR SYBR green mix (Eurogentec, Seraing, Belgium) for detection according to the manufacturer’s instructions. A standard curve was included on each plate by diluting genomic DNA from pure culture. *Dysosmobacter* spp standard curve ranged between 3.2 10^3^ and 1.0 10^7^ CFU per well for *Dysosmobacter* spp and between 6.4 10^2^ and 2.0 10^6^ CFU per well for total bacteria *(based on L. acidophilus DSM20079)*.

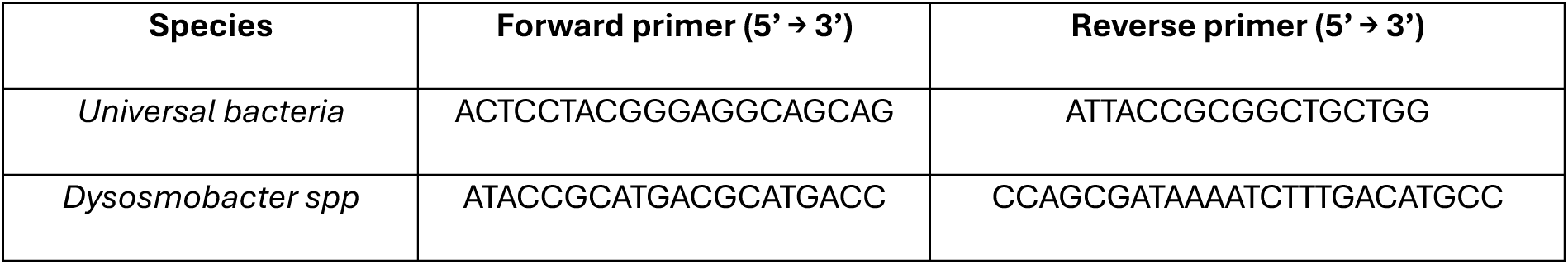

### Statistical analysis

Spearman’s correlation of *Dysosmobacter spp.* per gram of feces against all metabolites in each dataset was computed on RStudio (RStudio, 2023, Posit team) using the package ‘Psych’ (version 2.3.6) (24). A multiple testing correction via false discovery rate (FDR) estimation according to the Benjamini and Hochberg procedure was applied. The metabolites that significantly correlated with *Dysosmobacter spp.* (adjusted p-value <0.05) concentration in the feces after correction were extracted and listed in a table format ***(Table 2)***. Correlograms representing the correlations between *Dysosmobacter spp* fecal concentration and metabolites subfamilies in which one or multiple metabolites were associated were generated using the R package ‘Corrplot’ (version 0.92) (25). In these correlograms, while selected subfamilies were represented separately for clearer representation, the correlation scores and adjusted p-value represented are those obtained during the computation of the correlation between all 453 lipids and *Dysosmobacter spp*.

These metabolites were then used to explore correlation with metabolic and inflammatory parameters (hsCRP, IL-6, WBC, MCP-1, sCD40L, eotaxin, MCD, IP-10, Urea, Creatinine, GFR, sensitivity and resistance score, GIP, leptin, glucagon, BW, waist and hip circumference, waist-hip ratio, total cholesterol, LDL, HDL, total cholesterol/HDL ratio, TG, NEFAs, AST, ALT, GGT, AlkP, CK, LDH, iFABP, DDP4 activity, GLP-1, GLP-2, PYY, fasting glycemia, insulinemia, HbA1c proportion, BMI, fat mass, lean mass, diastolic and systolic blood pressure). The significant associations after FDR correction were listed in a table ***(Table 3)***.

### Abbreviations

In the correlograms for readability, the names of some lipid species have been shortened. DAG, PC, lysoPC, PE, lysoPE, PI and lysoPI names were shortened to only display the acyl chains composition. Myristoyl-linoleoyl-glycerol (14:0/18:2) [1]* = DAG(14:0/18:2) [1]*, palmitoyl-oleoyl-glycerol (16:0/18:1) [1]* = DAG(16:0/18:1) [1]*, palmitoyl-oleoyl-glycerol (16:0/18:1) [2]* = DAG (16:0/18:1) [2]*, palmitoyl-linoleoyl-glycerol (16:0/18:2) [1]* = DAG(16:0/18:2) [1]*, palmitoyl-linoleoyl-glycerol (16:0/18:2) [2]* = DAG (16:0/18:2) [2]*, palmitoyl-linolenoyl-glycerol (16:0/18:3) [2]* = DAG(16:0/18:3) [2]*, palmitoleoyl-oleoyl-glycerol (16:1/18:1) [1]* = DAG (16:1/18:1) [1]*, palmitoleoyl-linoleoyl-glycerol (16:1/18:2) [1]* = DAG (16:1/18:2) [1]*, palmitoyl-arachidonoyl-glycerol (16:0/20:4) [1]* = DAG(16:0/20:4) [1]*, palmitoyl-arachidonoyl-glycerol (16:0/20:4) [2]* = DAG(16:0/20:4) [2]*, oleoyl-oleoyl-glycerol (18:1/18:1) [1]* = DAG(18:1/18:1) [1]*, oleoyl-oleoyl-glycerol (18:1/18:1) [2]* = DAG(18:1/18:1) [2]*, oleoyl-linoleoyl-glycerol (18:1/18:2) [1] = DAG (18:1/18:2) [1], oleoyl-linoleoyl-glycerol (18:1/18:2) [1] = DAG (18:1/18:2) [2], oleoyl-linolenoyl-glycerol (18:1/18:3) [2]* = DAG (18:1/18:3) [2]*, linoleoyl-linoleoyl-glycerol (18:2/18:2) [1]* = DAG (18:2/18:2) [1]*, linoleoyl-linoleoyl-glycerol (18:2/18:2) [2]* = DAG (18:2/18:2) [2]*, linoleoyl-linolenoyl-glycerol (18:2/18:3) [1]* = DAG (18:2/18:3) [1]*, linoleoyl-linolenoyl-glycerol (18:2/18:3) [2]* = DAG (18:2/18:3) [2]*, stearoyl-arachidonoyl-glycerol (18:0/20:4) [1]* = DAG (18:0/20:4) [1]*, stearoyl-arachidonoyl-glycerol (18:0/20:4) [2]* = DAG (18:0/20:4) [2]*, oleoyl-arachidonoyl-glycerol (18:1/20:4) [1]* = DAG (18:1/20:4) [1]*, oleoyl-arachidonoyl-glycerol (18:1/20:4) [2]* = DAG (18:1/20:4) [2]*, linoleoyl-arachidonoyl-glycerol (18:2/20:4) [1]* = DAG (18:2/20:4) [1]*, linoleoyl-arachidonoyl-glycerol (18:2/20:4) [2]* = DAG (18:2/20:4) [2]*, linoleoyl-docosahexaenoyl-glycerol (18:2/22:6) [2]* = DAG (18:2/22:6) [2]*, 1-myristoyl-2-palmitoyl-GPC (14:0/16:0) = PC (14:0/16:0), 1-myristoyl-2-arachidonoyl-GPC (14:0/20:4)* = PC (14:0/20:4)*, 1,2-dipalmitoyl-GPC (16:0/16:0) = PC (16:0/16:0), 1-palmitoyl-2-palmitoleoyl-GPC (16:0/16:1)* = PC (16:0/16:1)*, 1-palmitoyl-2-stearoyl-GPC (16:0/18:0) = PC (16:0/18:0), 1-palmitoyl-2-oleoyl-GPC (16:0/18:1) = PC (16:0/18:1), 1-palmitoyl-2-linoleoyl-GPC (16:0/18:2) = PC (16:0/18:2), 1-palmitoyl-2-alpha-linolenoyl-GPC (16:0/18:3n3)* = PC (16:0/18:3n3)*, 1-palmitoyl-2-gamma-linolenoyl-GPC (16:0/18:3n6)* = PC (16:0/18:3n6)*, 1-palmitoyl-2-dihomo-linolenoyl-GPC (16:0/20:3n3 or 6)* = PC (16:0/20:3n3 or 6)*, 1-palmitoyl-2-arachidonoyl-GPC (16:0/20:4n6) = PC (16:0/20:4n6), 1-palmitoyl-2-docosahexaenoyl-GPC (16:0/22:6) = PC (16:0/22:6), 1-palmitoleoyl-2-linoleoyl-GPC (16:1/18:2)* = PC (16:1/18:2)*, 1-palmitoleoyl-2-linolenoyl-GPC (16:1/18:3)* = PC (16:1/18:3)*, 1,2-distearoyl-GPC (18:0/18:0) = PC (18:0/18:0), 1-stearoyl-2-oleoyl-GPC (18:0/18:1) = PC (18:0/18:1), 1-stearoyl-2-linoleoyl-GPC (18:0/18:2)* = PC (18:0/18:2)*, 1-stearoyl-2-arachidonoyl-GPC (18:0/20:4) = PC (18:0/20:4), 1-stearoyl-2-docosahexaenoyl-GPC (18:0/22:6) = PC (18:0/22:6), 1-oleoyl-2-linoleoyl-GPC (18:1/18:2)* = PC (18:1/18:2)*, 1-oleoyl-2-docosahexaenoyl-GPC (18:1/22:6)* = PC (18:1/22:6)*, 1,2-dilinoleoyl-GPC (18:2/18:2) = PC (18:2/18:2), 1-linoleoyl-2-linolenoyl-GPC (18:2/18:3)* = PC (18:2/18:3)*, 1-linoleoyl-2-arachidonoyl-GPC (18:2/20:4n6)* = PC (18:2/20:4n6)*, 1-palmitoyl-GPC (16:0) = PC (16:0), 2-palmitoyl-GPC* (16:0)* = PC* (16:0)*, 1-palmitoleoyl-GPC* (16:1)* = PC* (16:1)* [1], 2-palmitoleoyl-GPC* (16:1)* = PC* (16:1)* [2], 1-oleoyl-GPC (18:1) = PC (18:1), 1-linoleoyl-GPC (18:2) = PC (18:2), 1-linoleoyl-GPC (18:3)* = PC (18:3)*, 1-arachidonoyl-GPC* (20:4)* = PC* (20:4)*, 1-lignoceroyl-GPC (24:0) = PC (24:0), 1-palmitoyl-2-oleoyl-GPE (16:0/18:1) = PE (16:0/18:1), 1-palmitoyl-2-linoleoyl-GPE (16:0/18:2) = PE (16:0/18:2), 1-palmitoyl-2-arachidonoyl-GPE (16:0/20:4)* = PE (16:0/20:4)*, 1-palmitoyl-2-docosahexaenoyl-GPE (16:0/22:6)* = PE (16:0/22:6)*, 1-stearoyl-2-oleoyl-GPE (18:0/18:1) = PE (18:0/18:1), 1-stearoyl-2-linoleoyl-GPE stearoyl-2-docosahexaenoyl-GPE (18:0/22:6)* = PE (18:0/22:6)*, 1-oleoyl-2-linoleoyl-GPE (18:1/18:2)* = PE (18:1/18:2)*, 1-oleoyl-2-arachidonoyl-GPE (18:1/20:4)* = PE (18:1/20:4)*, 1-oleoyl-2-docosahexaenoyl-GPE (18:1/22:6)* = PE (18:1/22:6)*, 1,2-dilinoleoyl-GPE (18:2/18:2)* = PE (18:2/18:2)*, 1-palmitoyl-GPE (16:0) = PE (16:0), 1-stearoyl-GPE (18:0) = PE (18:0) [1], 2-stearoyl-GPE (18:0)* [2] = PE (18:0)*, 1-oleoyl-GPE (18:1) = PE (18:1), 1-linoleoyl-GPE (18:2)* = PE (18:2)*, 1-palmitoyl-2-oleoyl-GPI (16:0/18:1)* = PI (16:0/18:1)*, 1-palmitoyl-2-linoleoyl-GPI (16:0/18:2) = PI (16:0/18:2), 1-palmitoyl-2-arachidonoyl-GPI (16:0/20:4)* = PI (16:0/20:4)*, 1-stearoyl-2-oleoyl-GPI (18:0/18:1)* = PI (18:0/18:1)*, 1-stearoyl-2-linoleoyl-GPI (18:0/18:2) = PI (18:0/18:2), 1-stearoyl-2-arachidonoyl-GPI (18:0/20:4) = PI (18:0/20:4), 1-palmitoyl-GPI* (16:0) = PI* (16:0), 1-stearoyl-GPI (18:0) = PI (18:0), 1-oleoyl-GPI (18:1) = PI (18:1), 1-linoleoyl-GPI* (18:2)* = PI* (18:2)*, 1-arachidonoyl-GPI* (20:4)* = PI* (20:4)*.

## Results

In a previous study in mice, supplementation with *D. welbionis* J115^T^ under high-fat diet (60%, HFD) altered the lipid profile in the plasma, the brown adipose tissue (BAT) and the colon. Using a targeted lipidomics approach focused on a small subset of lipids (65 targeted), we observed that in plasma, six lipids (PGD2, 17-HDoHE, C18-3OH, C14-3OH, C18-2OH and 12-HETE) were restored to levels comparable to the control group, while two lipids (15-dPGJ2, C16-3OH) were decreased. Additionally, 14,15-EET was elevated compared to normal diet (ND), 9oxoODE was decreased compared to HFD, and three lipids (13-oxoODE, 9-HODE and 10-HODE) were decreased compared to both ND and untreated HFD groups (26).

Building on these findings, we extended our investigation to humans by analyzing the association of *Dysosmobacter spp.* with plasmatic lipid metabolites in the Microbes4U cohort. This study included 30 women and 22 men who were overweight or obese based on BMI and had a metabolic syndrome and insulin resistance ***(Table 1)***. Metabolomic profiling detected a total of 1169 compounds, 935 of which were of known identity, while 234 were of unknown identity (X-number) or partially characterized. Because of a very weak detection rate, metabolites related to drugs (i.e. analgesic, neurological, respiratory, antibiotic, psychoactive) and tobacco were not considered in the analysis, which gave us a final dataset of 862 metabolites. Based on Metabolon’s super-pathway classification, the metabolites were split into two datasets. The first one containing the 453 metabolites referred as belonging to the “Lipid” super-pathway, and the second one, “non-lipid”, encompassing the remaining 405 metabolites.

**Table 1:** Participants characteristics.

|  | Mean ±SEM |
| --- | --- |
| **Gender (nb(%))** |  |
| Women | 30 (57.69) |
| Men | 22 (42.30) |
| **Age (years)** | 51.46 ± 1.096 |
| **Height (cm)** | 1.71 ± 0.013 |
| **BMI** | 37.95 ± 0.681 |
| **Waist (cm)** | 122.59 ± 1.657 |
| **Hip (cm)** | 120.52 ± 1.577 |
| **Waist-hip ratio** | 1.02 ± 0.013 |
| **Fat mass (Kg)** | 46.02 ± 1.59 |
| **Lean mass (Kg)** | 65.79 ± 1.944 |
| **Systolic blood pressure (mmHg)** | 144.09 ± 2.169 |
| **Diastolic blood pressure (mmHg)** | 95.48 ± 2.122 |
| **Fasting blood glucose (mg/dl)** | 102.54 ± 1.512 |
| **Glycated hemoglobin A1c (%)** | 5.7 ± 0.061 |
| **Insulinemia (pmol/L)** | 145.78 ± 9.854 |
| **Insulin sensitivity (%)** | 32.92 ± 2.397 |
| **Total Cholesterol (mg/dl)** | 202.06 ± 5.632 |
| **LDL cholesterol (mg/dl)** | 129.65 ± 4.778 |
| **HDL cholesterol (mg/dl)** | 44.08 ± 1.116 |
| **Total cholesterol/HDL ratio** | 4.67 ± 0.138 |
| **AST activity (U/L)** | 23.52 ± 2.35 |
| **ALT activity (U/L)** | 33.35 ± 3.234 |
| **GGT activity (U/L)** | 40.69 ± 4.747 |
| **LDH activity (UI/L)** | 183.58 ± 4.245 |
| **CK activity (U/L)** | 119.18 ± 9.268 |
| **TG (mg/dl)** | 141.65 ± 6.845 |
| **Body weight (Kg)** | 111.14 ± 2.551 |
| **DDP4 activity (mU/ml)** | 15.52 ± 0.489 |

No correlation was found between *Dysosmobacter spp* fecal concentration and plasma non-lipid metabolites. Within the lipid super-pathway, only 9-HODE, previously identified in the mouse study, was detected. However, due to the detection method’s limited resolution, it was annotated as 9-HODE/13-HODE, as the two isomers could not be distinguished. No significant correlation was observed between this lipid and *Dysosmobacter spp* fecal concentration. Interestingly, 46 other lipid molecules showed positive correlations with *Dysosmobacter spp* fecal levels ***(Table 2)***. These metabolites had an average correlation coefficient (r-score) of 0.456, ranging from a minimum correlation of 0.387 for 1-palmitoyl-GPE (16:0) to a maximum of 0.544 for 1-palmitoyl-GPC (16:0).

**Table 2:** Plasma lipids correlated with *Dysosmobacter spp* fecal concentration.

|  | r-score | p-value | Adjusted p-value |
| --- | --- | --- | --- |
| **Diacylglycerol** |  |  |  |
| Diacylglycerol (14:0/18:1, 16:0/16:1) [1]* | 0.39310 | 0.00394 | 0.04488 |
| Linoleoyl-arachidonoyl-glycerol (18:2/20:4) [1]* | 0.41945 | 0.00197 | 0.02739 |
| Linoleoyl-arachidonoyl-glycerol (18:2/20:4) [2]* | 0.51567 | 0.00009 | 0.00265 |
| Linoleoyl-linolenoyl-glycerol (18:2/18:3) [2]* | 0.46911 | 0.00045 | 0.00917 |
| Myristoyl-linoleoyl-glycerol (14:0/18:2) [1]* | 0.45500 | 0.00070 | 0.01282 |
| Oleoyl-arachidonoyl-glycerol (18:1/20:4) [1]* | 0.48877 | 0.00024 | 0.00558 |
| Oleoyl-arachidonoyl-glycerol (18:1/20:4) [2]* | 0.50423 | 0.00014 | 0.00367 |
| Palmitoyl-arachidonoyl-glycerol (16:0/20:4) [1]* | 0.43149 | 0.00140 | 0.02143 |
| Palmitoyl-arachidonoyl-glycerol (16:0/20:4) [2]* | 0.53916 | 0.00004 | 0.00130 |
| Palmitoyl-linoleoyl-glycerol (16:0/18:2) [2]* | 0.39042 | 0.00422 | 0.04701 |
| Stearoyl-arachidonoyl-glycerol (18:0/20:4) [1]* | 0.52275 | 0.00007 | 0.00215 |
| Stearoyl-arachidonoyl-glycerol (18:0/20:4) [2]* | 0.53663 | 0.00004 | 0.00141 |
| **Monoacylglycerol** |  |  |  |
| 1-arachidonoylglycerol (20:4) | 0.43166 | 0.00140 | 0.02136 |
| 1-palmitoleoylglycerol (16:1)* | 0.39231 | 0.00402 | 0.04547 |
| **Acylcholines** |  |  |  |
| Arachidonoylcholine | 0.40630 | 0.00280 | 0.03524 |
| Eicosapentaenoylcholine | 0.40596 | 0.00283 | 0.03549 |
| Oleoylcholine | 0.41168 | 0.00243 | 0.03182 |
| Palmitoleoylcholine | 0.45420 | 0.00072 | 0.01305 |
| Palmitoylcholine | 0.40195 | 0.00314 | 0.03829 |
| Stearoylcholine* | 0.40451 | 0.00294 | 0.03647 |
| **Long Chain Polyunsaturated Fatty Acids** |  |  |  |
| Arachidonate (20:4n6) | 0.40203 | 0.00313 | 0.03823 |
| **Phosphatidylcholine** |  |  |  |
| 1-linoleoyl-2-arachidonoyl-GPC (18:2/20:4n6)* | 0.45121 | 0.00079 | 0.01395 |
| 1-myristoyl-2-arachidonoyl-GPC (14:0/20:4)* | 0.43618 | 0.00123 | 0.01942 |
| 1-palmitoyl-2-gamma-linolenoyl-GPC (16:0/18:3n6)* | 0.46779 | 0.00047 | 0.00948 |
| 1-palmitoyl-2-oleoyl-GPC (16:0/18:1) | 0.44404 | 0.00098 | 0.01636 |
| 1-stearoyl-2-oleoyl-GPC (18:0/18:1) | 0.42679 | 0.00160 | 0.02361 |
| **Lysophosphatidylcholine** |  |  |  |
| 1-lignoceroyl-GPC (24:0) | 0.40357 | 0.00301 | 0.03715 |
| 1-linolenoyl-GPC (18:3)* | 0.50645 | 0.00013 | 0.00344 |
| 1-oleoyl-GPC (18:1) | 0.48596 | 0.00026 | 0.00600 |
| 1-palmitoleoyl-GPC* (16:1)* | 0.53436 | 0.00004 | 0.00151 |
| 1-palmitoyl-GPC (16:0) | 0.54418 | 0.00003 | 0.00110 |
| 1-stearoyl-GPC (18:0) | 0.47887 | 0.00033 | 0.00721 |
| 2-palmitoleoyl-GPC* (16:1)* | 0.49142 | 0.00022 | 0.00521 |
| 2-palmitoyl-GPC* (16:0)* | 0.51900 | 0.00008 | 0.00241 |
| **Lysophosphatidylethanolamine** |  |  |  |
| 1-palmitoyl-GPE (16:0) | 0.38709 | 0.00458 | 0.04988 |
| 1-stearoyl-GPE (18:0) | 0.49244 | 0.00021 | 0.00507 |
| 2-stearoyl-GPE (18:0)* | 0.48707 | 0.00025 | 0.00583 |
| **Phosphatidylinositol** |  |  |  |
| 1-stearoyl-2-arachidonoyl-GPI (18:0/20:4) | 0.44574 | 0.00093 | 0.01573 |
| **Lysophosphatidylinositol** |  |  |  |
| 1-arachidonoyl-GPI* (20:4)* | 0.39076 | 0.00418 | 0.04671 |
| 1-palmitoyl-GPI* (16:0) | 0.46991 | 0.00044 | 0.00899 |
| 1-stearoyl-GPI (18:0) | 0.40348 | 0.00302 | 0.03722 |
| **Lysoplasmalogen** |  |  |  |
| GPC(P-16:0) | 0.42995 | 0.00147 | 0.02211 |
| **Phospholipid Metabolism** |  |  |  |
| Glycerophosphoethanolamine | 0.49825 | 0.00017 | 0.00433 |
| Glycerophosphorylcholine (GPC) | 0.46376 | 0.00053 | 0.01042 |
| **Ceramides** |  |  |  |
| Ceramide (d18:1/20:0, d16:1/22:0, d20:1/18:0)* | 0.51225 | 0.00010 | 0.00291 |
| **Sterol** |  |  |  |
| Cholesterol | 0.45087 | 0.00080 | 0.01406 |

\*Indicates a compound that has not been confirmed based on a standard, but that is identified by Metabolon with confidence. [#] Indicates a compound that is a structural isomer of another compound in the Metabolon spectral library

Since this study focused on plasmatic metabolites, all lipids detected were either free fatty acid derived from triglycerides lipolysis, circulating primarily bound to albumin, or from chylomicrons lysed during the sample preparation (27).

### *Dysosmobacter spp* partially correlates with DAGs and MAGs

Among the 28 measured diacylglycerols (DAGs), 12 showed a positive correlation with *Dysosmobacter spp* fecal concentrations, all of which contained at least one mono- or polyunsaturated acyl chain **(*Figure 1*)**. These included: diacylglycerol (14:0/18:1, 16:0/16:1) [1]*, myristoyl-linoleoyl-glycerol (14:0/18:2) [1]*, linoleoyl-linolenoyl-glycerol(18:2/18:3) [2]*, palmitoyl-linoleoyl-glycerol (16:0/18:2) [2]*, two isomers of palmitoyl-arachidonoyl-glycerol (16:0/20:4), two isomers of stearoyl-arachidonoyl-glycerol (18:0/20:4), two isomers of oleoyl-arachidonoyl-glycerol (18:1/20:4), and two isomers of linoleoyl-arachidonoyl-glycerol (18:2/20:4) ***(Table 2)***.

**Figure 1:**
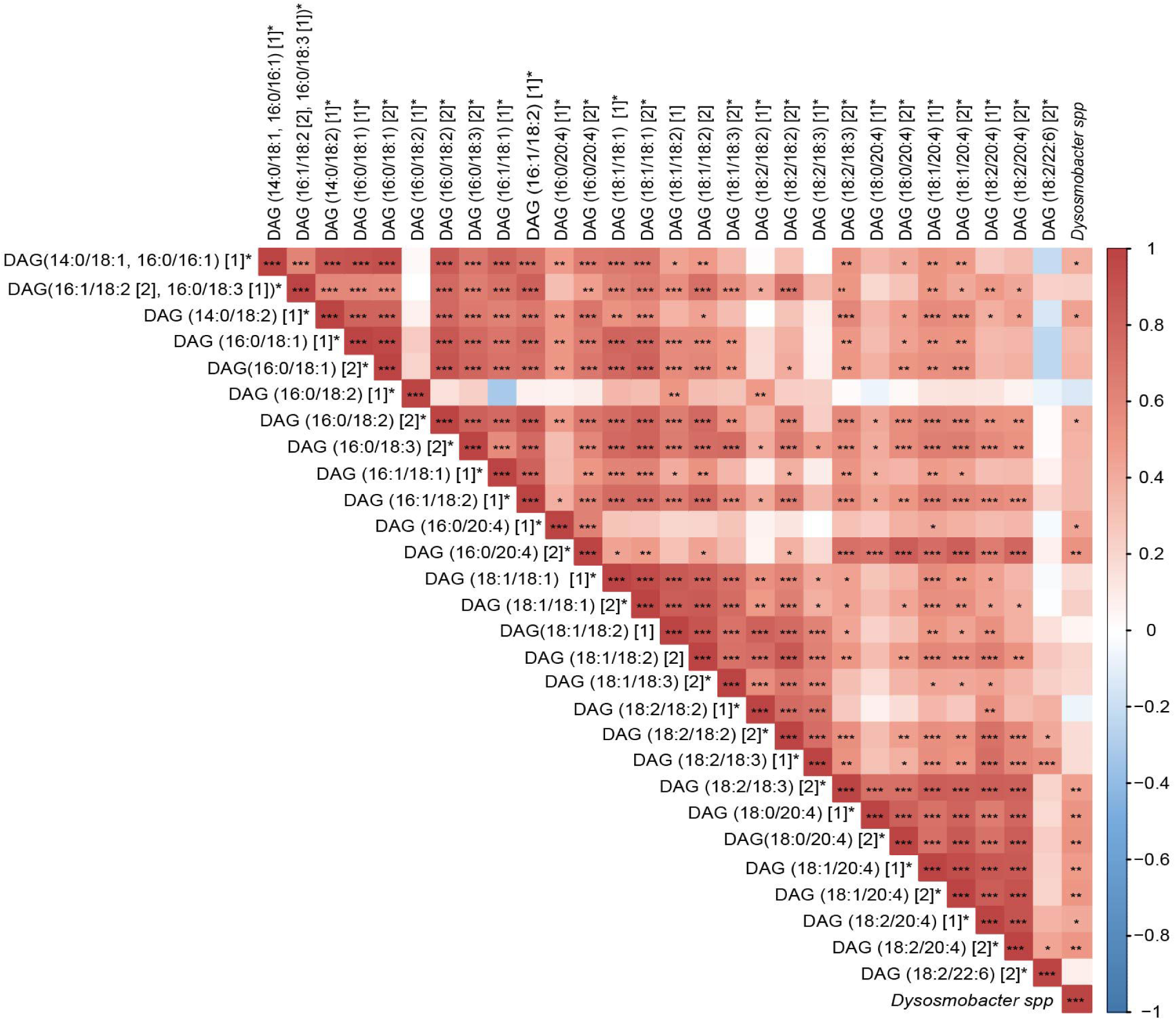
Spearman correlations between fecal Dysosmobacter spp and diacylglycerols. Positive correlations are labeled in red and negative ones in blue. Significant correlations after FDR correction are labeled as follow, *: adjusted-p <0.05, **: adjusted-p <0.01, ***: adjusted-p <0.001.

Notably, several of these DAGs were negatively correlated with GLP-1, an incretin involved in energy metabolism (28). Specifically, linoleoyl-arachidonoyl-glycerol (18:2/20:4) [1]*, palmitoyl-arachidonoyl-glycerol (16:0/20:4) [2]*, linoleoyl-linolenoyl-glycerol (18:2/18:3) [2]*, and stearoyl-arachidonoyl-glycerol (18:0/20:4) [1] *. Furthermore, Palmitoyl-arachidonoyl-glycerol (16:0/20:4) [1]* was negatively associated with plasma LPS concentration and Stearoyl-arachidonoyl-glycerol (18:0/20:4) [2]* was positively associated with GDF-15 ***(Table 3)***.

**Table 3:** Identified lipids correlate with metabolic & inflammatory plasmatic parameters.

|  | r-score | p-value | Adjusted p-value |
| --- | --- | --- | --- |
| **GLP-1** |  |  |  |
| 1-arachidonoyl-GPI* (20:4)* | -0.37585 | 0.00715 | 0.03729 |
| Arachidonoylcholine | -0.41330 | 0.00285 | 0.01724 |
| Glycerophosphoethanolamine | -0.38343 | 0.00598 | 0.03213 |
| Glycerophosphorylcholine (GPC) | -0.36874 | 0.00841 | 0.04289 |
| Linoleoyl-arachidonoyl-glycerol (18:2/20:4) [1]* | -0.41263 | 0.00290 | 0.01752 |
| Linoleoyl-linolenoyl-glycerol (18:2/18:3) [2]* | -0.41532 | 0.00271 | 0.01651 |
| Oleoylcholine | -0.36749 | 0.00865 | 0.04403 |
| Palmitoleoylcholine | -0.40860 | 0.00322 | 0.01900 |
| Palmitoyl-arachidonoyl-glycerol (16:0/20:4) [2]* | -0.36913 | 0.00834 | 0.04255 |
| Palmitoylcholine | -0.36634 | 0.00888 | 0.04482 |
| Stearoyl-arachidonoyl-glycerol (18:0/20:4) [1]* | -0.40994 | 0.00311 | 0.01852 |
| Stearoylcholine* | -0.46689 | 0.00063 | 0.00475 |
| **LPS** |  |  |  |
| 1-arachidonoyl-GPI* (20:4)* | -0.37742 | 0.00581 | 0.03144 |
| 1-palmitoyl-GPI* (16:0) | -0.44878 | 0.00085 | 0.00614 |
| 1-stearoyl-GPI (18:0) | -0.39604 | 0.00366 | 0.02125 |
| Palmitoyl-arachidonoyl-glycerol (16:0/20:4) [1]* | -0.38971 | 0.00429 | 0.02434 |
| **GDF-15** |  |  |  |
| 1-lignoceroyl-GPC (24:0) | 0.35976 | 0.00880 | 0.04453 |
| Stearoyl-arachidonoyl-glycerol (18:0/20:4) [2]* | 0.37574 | 0.00605 | 0.03238 |
| **IL-6** |  |  |  |
| 1-linolenoyl-GPC (18:3)* | -0.42002 | 0.00239 | 0.01484 |
| **MCP-1** |  |  |  |
| 1-oleoyl-GPC (18:1) | 0.36948 | 0.00702 | 0.03676 |
| 1-palmitoleoyl-GPC* (16:1)* | 0.35812 | 0.00914 | 0.04592 |
| 1-stearoyl-2-arachidonoyl-GPI (18:0/20:4) | 0.37217 | 0.00659 | 0.03489 |
| 1-stearoyl-2-oleoyl-GPC (18:0/18:1) | 0.42442 | 0.00171 | 0.01109 |
| 1-stearoyl-GPC (18:0) | 0.37708 | 0.00586 | 0.03159 |
| **LBP** |  |  |  |
| 1-palmitoleoyl-GPC* (16:1)* | 0.37096 | 0.00678 | 0.03568 |
| 1-palmitoyl-2-oleoyl-GPC (16:0/18:1) | 0.43960 | 0.00111 | 0.00765 |
| 1-stearoyl-2-oleoyl-GPC (18:0/18:1) | 0.43319 | 0.00134 | 0.00893 |
| 2-palmitoleoyl-GPC* (16:1)* | 0.38043 | 0.00540 | 0.02957 |
| **AlkP** |  |  |  |
| 1-palmitoyl-GPC (16:0) | 0.37648 | 0.00647 | 0.03433 |
| 1-stearoyl-GPE (18:0) | 0.38848 | 0.00485 | 0.02711 |
| **Waist-hip ratio** |  |  |  |
| 1-stearoyl-2-arachidonoyl-GPI (18:0/20:4) | 0.4108 | 0.00228 | 0.01422 |
| **HbA1c** |  |  |  |
| 2-palmitoleoyl-GPC* (16:1)* | -0.35592 | 0.00961 | 0.04814 |
| **Urea** |  |  |  |
| Arachidonate (20:4n6) | -0.36366 | 0.00805 | 0.04128 |
| **ADAMts13** |  |  |  |
| GPC(P-16:0) | 0.46398 | 0.00053 | 0.00410 |

\*Indicates a compound that has not been confirmed based on a standard, but that is identified by Metabolon with confidence
[#] Indicates a compound that is a structural isomer of another compound in the Metabolon spectral library

In general, several plasmatic DAGs have been positively linked to insulin resistance, T2D progression, and related markers in various cohorts, including non-diabetic overweight/obese individuals (29), non-diabetic individuals with high risk of cardiovascular disease (CVD), and those with T2D (30). However, in our cohort of insulin resistant but non-diabetic participants, DAGs showed no correlation with key glucose metabolism parameters, including fasting glycemia, HbA1c, insulin sensitivity and, insulin resistance scores.

One possible explanation for these discrepancies is the variation in how DAG species are reported across studies. Like for many lipid families, many studies aggregate DAGs as a total lipid class or sum the carbon and double-bound content of their acyl chains, rendering direct comparison between studies difficult. The specific composition and localization of DAG species may influence their metabolic effects, and these differences should be considered in future investigations.

Among the monoacylglycerols (MAGs), 1-palmitoleoylglycerol (16:1)* and 1-arachidonylglycerol (20:4) (1-AG) were both positively correlated with *Dysosmobacter spp* fecal concentration (r =0.392, adjusted p-value = 0.045; and r=0.431, adjusted p-value =0.021 respectively) ***(Table 1 and* *Figure 2*)**. MAGs are typically produced from DAGs via diacylglycerol lipases, with 2-monoacylglycerols (2-MAGs) capable of non-enzymatic isomerization into 1/3-MAGs, which are thermodynamically more stable (31). This may occur *in vivo* or during the analytical process. The analytical method used in this study does not distinguish between 1- and 3-MAG isomers, so these molecules are collectively labeled as 1-MAGs.

**Figure 2:**
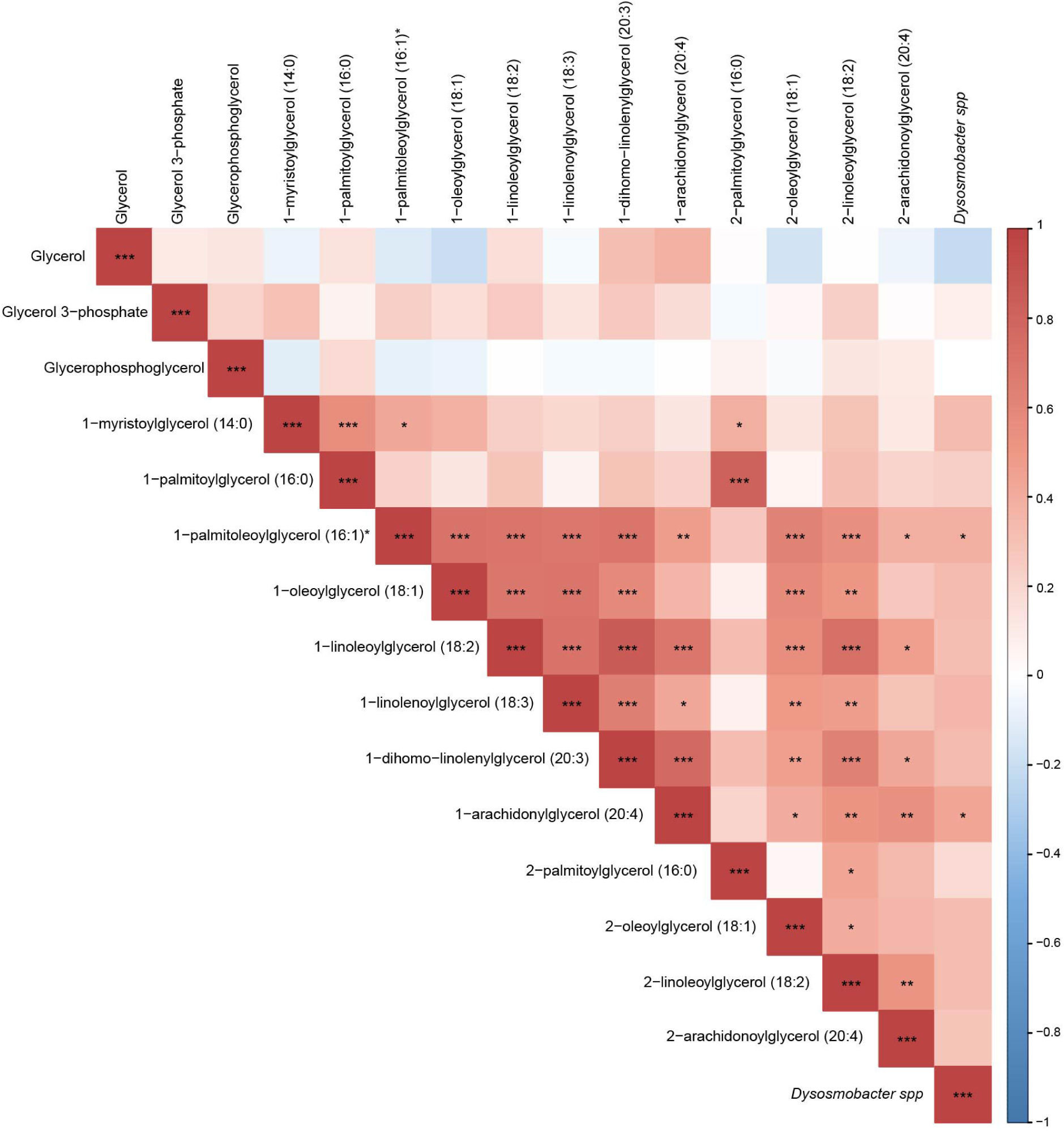
Spearman correlations between fecal Dysosmobacter spp and monoacylglycerols. Positive correlations are labeled in red and negative ones in blue. Significant correlations after FDR correction are labeled as follow, *: adjusted-p <0.05, **: adjusted-p <0.01, ***: adjusted-p <0.001.

Very few studies have reported the detection of plasmatic 1-palmitoleoylglycerol and 1-AG in human cohorts focused on metabolic diseases. In a Finnish male cohort, plasma 1-palmitoleoylglycerol was positively associated with T2D incidence over a seven-year follow-up (32). Similarly, in a 2022 study of Qatari individuals with T2D, 1-palmitoleoylglycerol was linked to higher BMI, elevated triglycerides levels, dyslipidemia and lower HDL levels (33). Mechanistically, an *in vitro* study conducted in 2017 suggests that 1-AG could participate in the stabilization and maintenance of the cannabinoid receptor 1 (CB1), a receptor involved, among other roles, in increased intestinal permeability – a feature which may contribute to metabolic syndrome (34, 35). In our cohort, these 2 metabolites did not correlate with any of the metabolic or inflammatory parameters measured in the participants.

Despite the typical association of DAGs and MAGs with metabolic dysfunction, 14 of these were associated with *Dysosmobacter spp* fecal concentration in our study. Given that *Dysosmobacter spp.* has been linked to beneficial metabolic outcomes, this unexpected finding raises important questions about its role in lipid metabolism. Future studies are needed to elucidate whether these correlations reflect a protective mechanism, a compensatory response, or an unrelated phenomenon.

### Dysosmobacter spp correlates with acylcholines

Fecal *Dysosmobacter spp* concentration positively correlated with six of the nine plasma acylcholines detected: palmitoylcholine, oleoylcholine, palmitoloelycholine, stearoylcholine*, arachidonoylcholine and eicosapentaenoylcholine ***(Table 2 and* *Figure 3*)**. Notably, all identified acylcholines -except eicosapentaenoylcholine-were negatively associated with plasma GLP-1 plasmatic levels ***(Table 3)***, suggesting a potential metabolic interaction.

**Figure 3:**
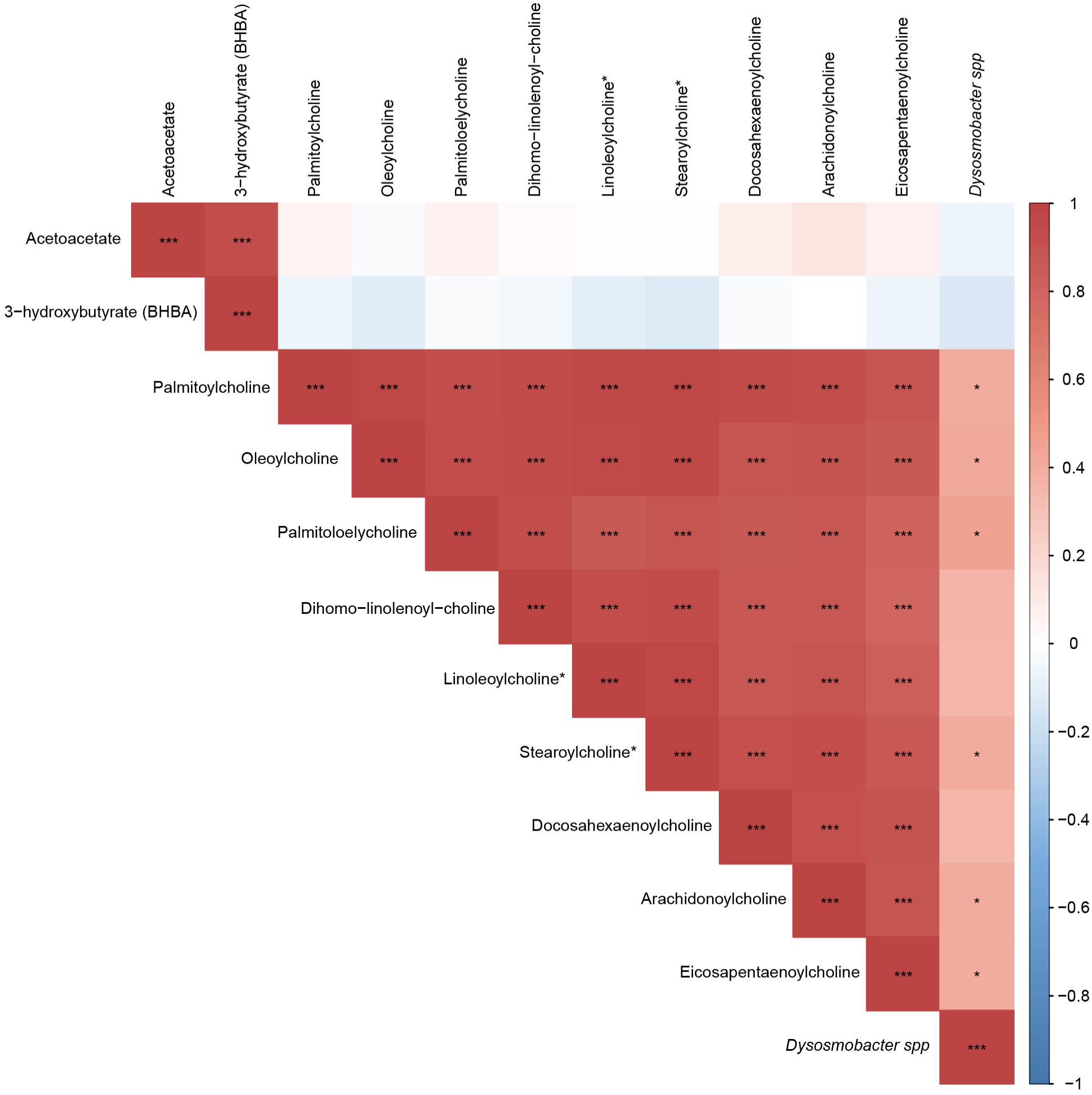
Spearman correlations between fecal Dysosmobacter spp and acylcholines. Positive correlations are labeled in red and negative ones in blue. Significant correlations after FDR correction are labeled as follow, *: adjusted-p <0.05, **: adjusted-p <0.01, ***: adjusted-p <0.001.

**Figure 4:**
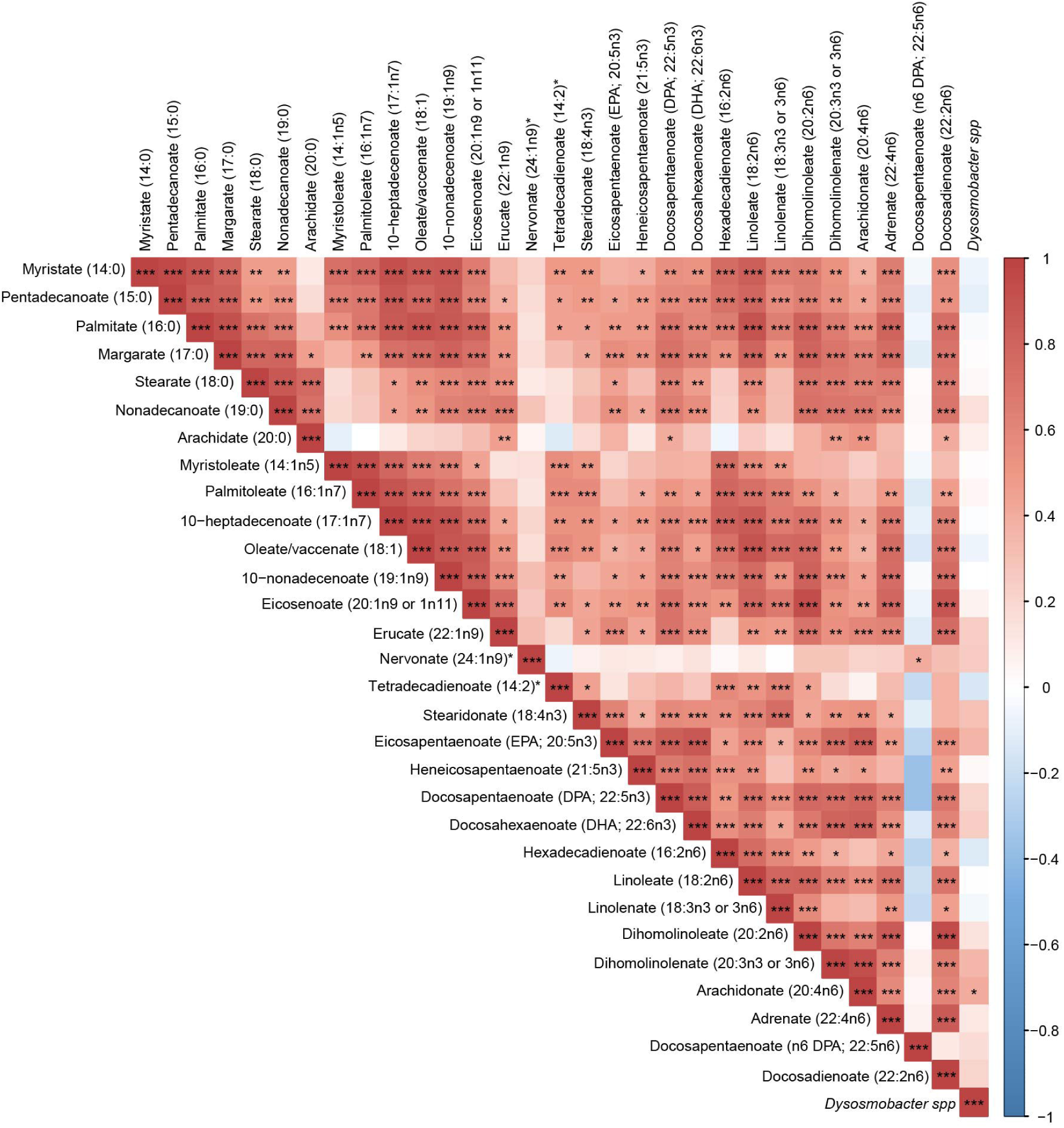
Spearman correlations between fecal Dysosmobacter spp and long chain polyunsaturated fatty acids. Positive correlations are labeled in red and negative ones in blue. Significant correlations after FDR correction are labeled as follow, *: adjusted-p <0.05, **: adjusted-p <0.01, ***: adjusted-p <0.001.

Despite their presence in circulation, the role of plasma acylcholines in metabolic syndrome remains largely unexplored. In a 2023 Qatari cohort of individuals with T2D treated with metformin, palmitoylcholine and arachidonoylcholine were associated with a better treatment response (36). This is particularly intriguing given that a previous study from our lab demonstrated an increased abundance of *Dysosmobacter spp* in metformin-treated diabetic individuals. However, in an experiment in which mice were co-treated with both metformin and *D. welbionis* J115^T^ no synergistic effect was observed.

Beyond metabolic syndrome, plasmatic acylcholines variations show inconsistent associations across different pathological contexts. They have been positively linked to endometrial cancer (37), pulmonary embolism risk (38) and atherosclerotic plaques (39), while negatively associated with chronic thromboembolic pulmonary hypertension (40) and myalgic encephalomyelitis/chronic fatigue syndrome (41). Additionally, in clinical trials where prebiotic interventions aimed to modulate gut microbiota, plasmatic acylcholine profile were altered (42–45), highlighting a potential microbiota-mediated influence on acylcholine metabolism. However, these changes were highly variable and dependent on cohort composition and the type of prebiotic used.

First described in 1911, long-chain acylcholines were initially studied for their pressor effect, conferring them a role in blood pressure regulation (46). Research on their biological activity declined after the 1950s, resulting in limited data (41, 46), but recent studies have reignited interest. In vitro experiments have shown that arachidonoylcylcholine, oleoylcholine, and linoleoylcholine, act as inhibitors of nicotinic acetylcholine receptors (nAChR), while arachidonoylcholine can also modestly inhibit acetylcholinesterase (AChE) and butyrylcholinesterase (BChE) at higher concentrations (47). While we don’t know the pathophysiological concentration range of these molecules in various tissues, these findings suggest that acylcholines may play a role in endogenous acetylcholine signaling.

BChE (previously pseudocholinesterase), a nonspecific cholinesterase enzyme synthesized in the liver and released into the plasma either in free form or bound to LDLs, has been implicated in metabolic disorders. Increased plasma BChE activity has been described in patients with hyperlipidemia (48), and correlates with metabolic syndrome markers in diabetic and non-diabetic individuals (49, 50). BChE also plays a role in the hydrolysis of octanoyl-ghrelin, an orexigenic hormone, converting it to its inactive form (51). However, the resulting des-acyl ghrelin may have cell-proliferative effects, potentially stimulating adipogenesis and cardiovascular alterations. This suggests a functional role for BChE in the development and progression of both obesity and coronary artery disease (50). The complexity of BChE’s role in energy metabolism is further underscored by mouse knockout models, where BChE deficiency led to increased obesity under a high-fat diet, indicating that the role of BChE in energy metabolism is still far from being understood.

While the gut microbiota appears to influence plasma acylcholine regulation, the underlying mechanisms remain poorly characterized. The observed link between *Dysosmobacter spp,* acylcholines, and metformin response warrants further exploration to clarify this association and its metabolic implications.

### Plasmatic Arachidonate correlates with *Dysosmobacter spp*

Plasma levels of arachidonate (20:4n6), showed a positive correlation with *Dysosmobacter spp fecal concentration (r*= 0.402, adjusted p-value = 0.038) ***(Table 2 and figure 4)***. Arachidonate is the second most mobilized fatty acid during fasting (52) and plays a central role in numerous physiological and pathological processes. However, its precise impact on metabolic diseases remains unclear (53, 54).

In this cohort, plasma arachidonate levels were negatively correlated with urea ***(Table 3)***, although urea levels remained within the physiological range for all participants based on Belgian clinical guidelines (55).

The role of arachidonate in metabolic disorders is complex and context-dependent. In T2D, increased free arachidonate has been linked to oxidative stress, a key driver of insulin resistance in adipocytes and muscle tissue, as well as impaired insulin secretion (53, 54). Conversely, other studies suggest an inverse correlation with T2D and a protective role. In a very limited cohort of fasting women, arachidonate was inversely correlated with glycemia, and in an another T2D cohort, individuals with lower circulatory arachidonate levels exhibited more pronounced diabetic characteristics, which were attenuated upon arachidonic acid supplementation (53, 56).

These discrepancies may stem from differences in arachidonate metabolism. While arachidonate itself has been shown to stimulate insulin secretion, its downstream metabolites exert divergent effects. For example, PGE2 contributes to pancreatic beta-cell dysfunction, whereas metabolites such as epoxyeicosatrienoic acids (EETs) and 20-hydroxyeicosatetraenoic acid (20-HETE) from the cytochrome P450 (CYP450) pathway, or lipoxin A4 (LXA4) from the lipoxygenase (LOX) pathway, have opposing, potentially beneficial effects (54, 57, 58).

Arachidonate is best known for its role in inflammation, a key component of the metabolic syndrome. However, as described above, its impact depends on its enzymatic conversion. Arachidonate can be processed by cyclooxygenases (COX), LOX or CYP450, producing a wide variety of metabolites that are either pro-inflammatory (e.g. PGE2, Leukotrienes, HETEs) or anti-inflammatory (e.g. LXA4) properties (53, 59).

Since our study did not include a complete profiling of arachidonate-derived metabolites, we cannot determine whether *Dysosmobacter spp* is associated with reduced arachidonate metabolism – potentially leading to a pro-inflammatory state - or whether it promotes a metabolic shift toward either pro- or anti-inflammatory lipid mediators. Further investigations are needed to clarify the implications of *Dysosmobacter spp.* in arachidonate metabolism and its potential role in metabolic health.

### Phospholipids, lysophospholipids & lysoplasmalogens correlations with *Dysosmobacter spp*

Phospholipids (PL) are amphipathic molecules that serve as the main structural components of cell membranes. They are composed of a glycerol backbone, a polar head group (e.g. choline, serine, inositol, ethanolamine, or glycerol), and two acyl chains. Hydrolysis of one acyl chains results in Lysophospholipids (LysoPL). If one of the fatty acid chains is attached to the backbone via a vinyl-ether bond instead of an ester bond, the PL classifies as a plasmenylphospholipids (also called a plasmalogen), while a LysoPL becomes a lysoplasmenylphospholpid (lysoplasmalogen). These molecules are further categorized based on their polar head group and acyl chain composition.

### Phosphocholines (PC) and lysophosphocholines (LysoPC)

Fecal *Dysosmobacter* spp concentrations correlated with five phosphocholines (PC), and eight lysophosphocholines (LysoPC) ***(Table 2 and* *Figure 5*)**. Notably, all five PC molecules contained at least one mono or poly-unsaturated acyl chain, which might be functionally relevant. Among them, 1-stearoyl-2-oleoyl-GPC (18:0/18:1) was positively associated with plasmatic monocyte chemoattractant protein-1 (MCP-1, CCL-2 in humans) (r-score = 0.424; adjusted p-value = 0.011), a marker of immune cell recruitment. This molecule, along with 1-palmitoyl-2-oleoyl-GPC (16:0/18:1) also correlated positively with lipopolysaccharide-binding protein (LBP) ***(Table 3)***, a marker of systemic endotoxemia.

**Figure 5:**
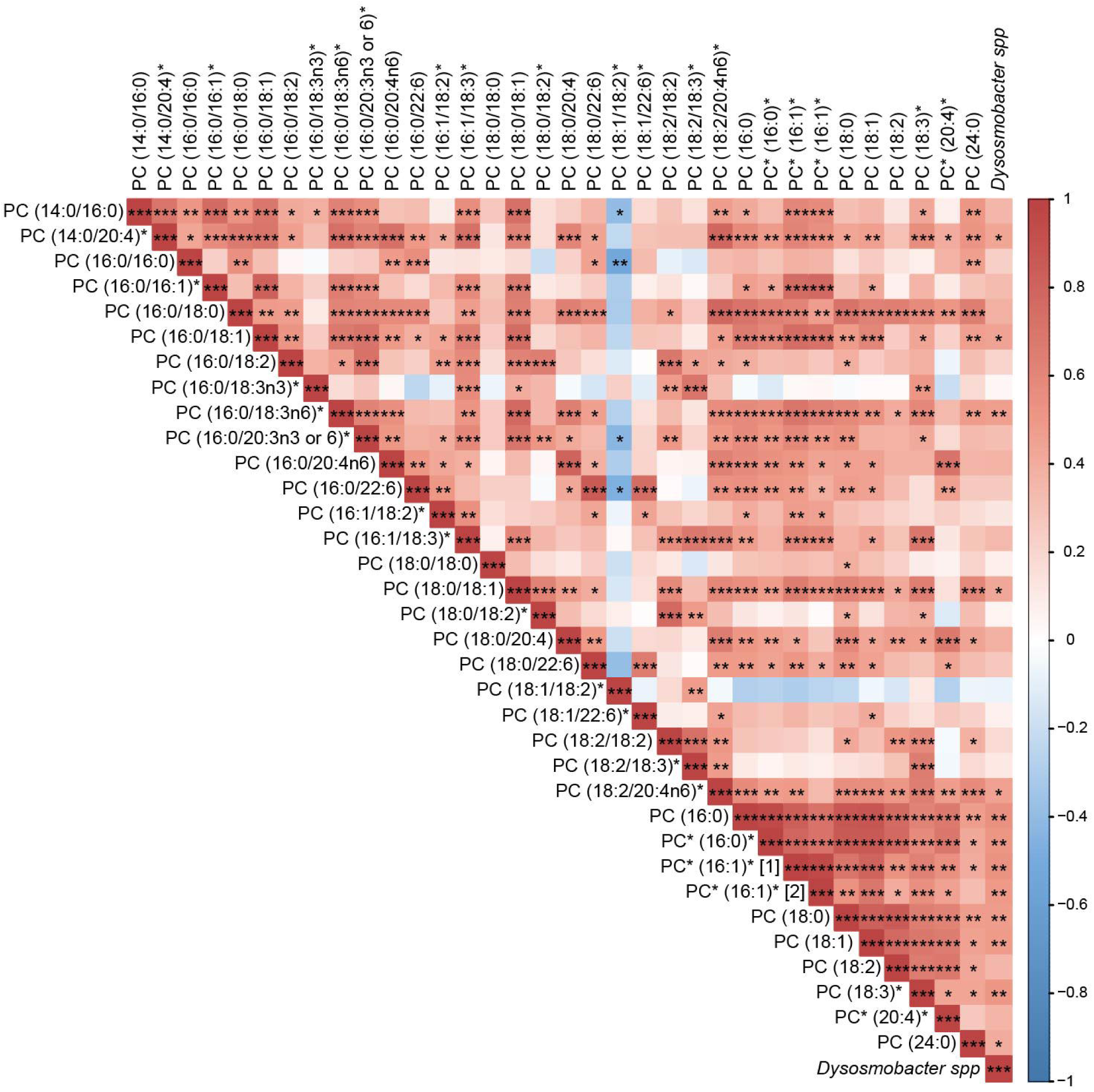
Spearman correlations between fecal Dysosmobacter spp, phosphatidylcholines and lysophosphatidylcholines. Positive correlations are labeled in red and negative ones in blue. Significant correlations after FDR correction are labeled as follow, *: adjusted-p <0.05, **: adjusted-p <0.01, ***: adjusted-p <0.001. For clarity on the correlogram, PC and LysoPC names were shortened to only display the acyl chains composition.

Previous studies have shown that the associations between circulating PC and metabolic syndrome vary widely and potentially depend on the differences in acyl chains length and saturation. A 2022 study carried out on Qatari cohort with and without T2D found that certain PC species correlated with BMI and dyslipidemia, but not with T2D or diabetic retinopathy (33). For example, PC (16:0/16:1) was positively associated with BMI and PC (16:0/18:0) was positively associated with LDL, triglycerides, LDL/HDL ratio and dyslipidemia, whereas PC (18:2/18:2) and PC (18:2/18:3) were negatively associated with BMI (33). In another cohort PC (16:0/20:4) was the only PC disrupted in prediabetic and diabetic patients, with elevated plasma levels (60). However, many studies reporting PC levels in metabolic disorders have lacked the analytical sensitivity to resolve individual molecular species, instead reporting summed acyl chain compositions, complicating cross-study comparisons (61–64).

Among the 10 detected LysoPCs, eight were positively correlated with *Dysosmobacter spp* fecal concentration ***(Table 2 and* *Figure 5*)**. Though present in lower quantities in cell membranes compared to their phospholipids counterparts (65–67), LysoPCs are abundant in human plasma, where their concentration range from 100 and 300µM in physiological state, with approximately 80% in the non-lipoprotein fraction, bound to albumin (29, 68). LysoPC are and have been implicated in the etiology of numerous disorders such as metabolic diseases, inflammation and cancer (67, 69).

In this study, 1-palmitoleoyl-GPC* (16:1)*, 1-oleoyl-GPC (18:1) and 1-stearoyl-GPC (18:0) were positively correlated positively with MCP-1 while 1-linolenoyl-GPC (18:3)* was negatively correlated with plasmatic IL-6. Both MCP-1 and IL-6 are markers of systemic inflammation and are often increased in obesity (70–75). 1-palmitoyl-GPC (16:0) correlated with plasmatic alkaline phosphatase (AlkP), a marker linked to hepatobiliary disorders and cardiometabolic risks (76, 77). 1-lignoceroyl GPC (24:0) positively correlated with plasmatic growth differentiation factor-15 (GDF-15), a member of the TGF-beta superfamily with complex effects on metabolism (78). Finally, both isomers of palmitoleoyl-GPC* (16:1)* were positively associated with plasma LBP levels, while one isomer was negatively associated with HbA1c, a marker reflecting long-term glycemic control ***(Table 3)***.

The relationship between LysoPCs and obesity appears to be highly dependent on acyl chain composition. Most human studies conducted have reported that plasma LysoPC (14:0) and LysoPC (16:1) are positively associated with obesity (61, 79, 80), whereas other lysoPCs tend to be negatively associated (68, 80–84) with a few exceptions (60, 85, 86). In this study, lysoPCs containing 16:0, 16:1, 18:0, 18:1, 18:3, and 24:0, were positively correlated with *Dysosmobacter spp*. but none correlated with weight parameters (BW, BMI, hip circumference, waist circumference) measured in our cohort.

Previous studies investigating the relationship between LysoPC and T2D, like in obesity, have predominantly reported an inverse correlation between plasma LysoPC levels and various T2D-related parameters. In particular, LysoPC species containing polyunsaturated fatty acids (PUFAs) tend to decrease with increasing insulin resistance (IR) and HOMA-IR scores (29, 62, 87, 88). Additionally, in two independent cohorts, baseline plasma LysoPC concentrations were

Despite these associations, the mechanisms underlying the potential beneficial effects of LysoPC on glucose metabolism remain unclear. While *in vitro* studies suggest that LysoPC can stimulate insulin secretion in pancreatic beta-cell lines further research is needed to fully understand this interaction (91).

In our study, none of the LysoPC correlated with *Dysosmobacter spp* fecal concentrations showed associations with glucose metabolism markers, including glucagon, insulin sensitivity, insulin resistance score, GLP-1 and leptin, excepted for the negative correlation between 2-palmitoleoyl-GPC* (16:1)*. When considering obesity and T2D comorbidities, research on LysoPC remains limited and often yields conflicting results. Some studies have reported negative associations between LysoPC levels hepatic fat accumulation, CVD risks, and cancer (29, 92–94), whereas findings on atherosclerosis and inflammation suggest both protective and detrimental roles depending on the context (69).

### Phosphatidylethanolamines (PE) and Lysophosphatidylethanolamines (LysoPE)

No correlations were found between *Dysosmobacter spp* fecal concentrations and total phosphatidylethanolamines (PE). However, three saturated LysoPE species - 2-stearoyl-GPE (18:0)* (r=0.487, adjusted p-value = 0.005), 1-stearoyl-GPE (18:0) (r=0.492, adjusted p-value = 0.005), and 1-palmitoyl-GPE (16:0) (r=0.387, adjusted p-value = 0.049) - were positively correlated ***(Table 2 and* *Figure 6*)**. These molecules have previously been identified as negatively associated with HOMA-IR in Chinese non-diabetic, non-obese individuals (62). While plasma PE levels are generally elevated in insulin-resistant individuals (60), LysoPEs are often reduced in overweight/obese participants (29, 86, 95), T2D patients, and inversely correlated with T2D and CVD risks (87, 94), and BMI (79). In this study, 1-stearoyl-GPE (18:0) was positively correlated with plasma AlkP ***(Table 3)***.

**Figure 6:**
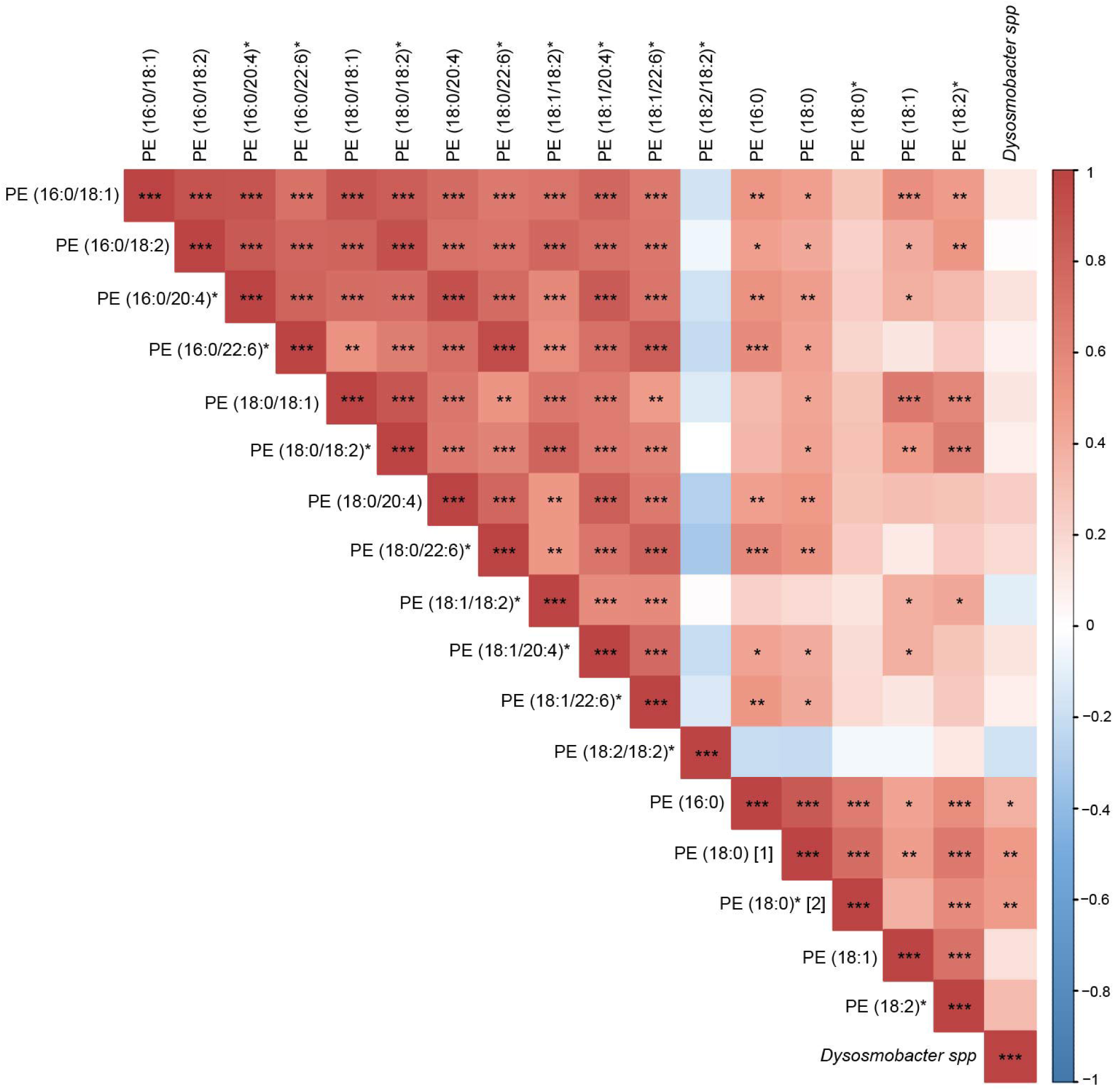
Spearman correlations between fecal Dysosmobacter spp, phosphatidylethanolamines and lysophosphatidylethanolamines. Positive correlations are as follow, *: adjusted-p <0.05, **: adjusted-p <0.01, ***: adjusted-p <0.001. For clarity on the correlogram, PE and LysoPE names were shortened to only display the acyl chains composition.

### Phosphatidylinositols (PI) and Lysophosphatidylinositols (LysoPI)

For Phosphatidylinositols (PI) and Lysophosphatidylinositols (LysoPI), *Dysosmobacter spp.* fecal concentrations correlated positively with plasma 1-stearoyl-2-arachidonoyl-GPI (18:0/20:4) (r-score= 0.445, adjusted p-value = 0.015), 1-palmitoyl-GPI* (16:0)) (r-score= 0.469, adjusted p-value = 0.008), 1-stearoyl-GPI (18:0) (r-score= 0.403, adjusted p-value = 0.037), and 1-arachidonoyl-GPI* (20:4)* (r-score= 0.390, adjusted p-value = 0.046) ***(Table 2 and* *Figure 7*)**. Among these, 1-arachidonoyl-GPI* (20:4)*, 1-palmitoyl-GPI* (16:0) and 1-stearoyl-GPI (18:0) were negatively associated with plasma LPS concentrations. In addition, 1-arachidonoyl-GPI* (20:4)* was negatively correlated with GLP-1, while 1-stearoyl-2-arachidonoyl-GPI (18:0/20/4) was positively associated with MCP-1 and waist-to-hip ratio, suggesting a potential link to visceral fat distribution ***(Table 3)***.

**Figure 7:**
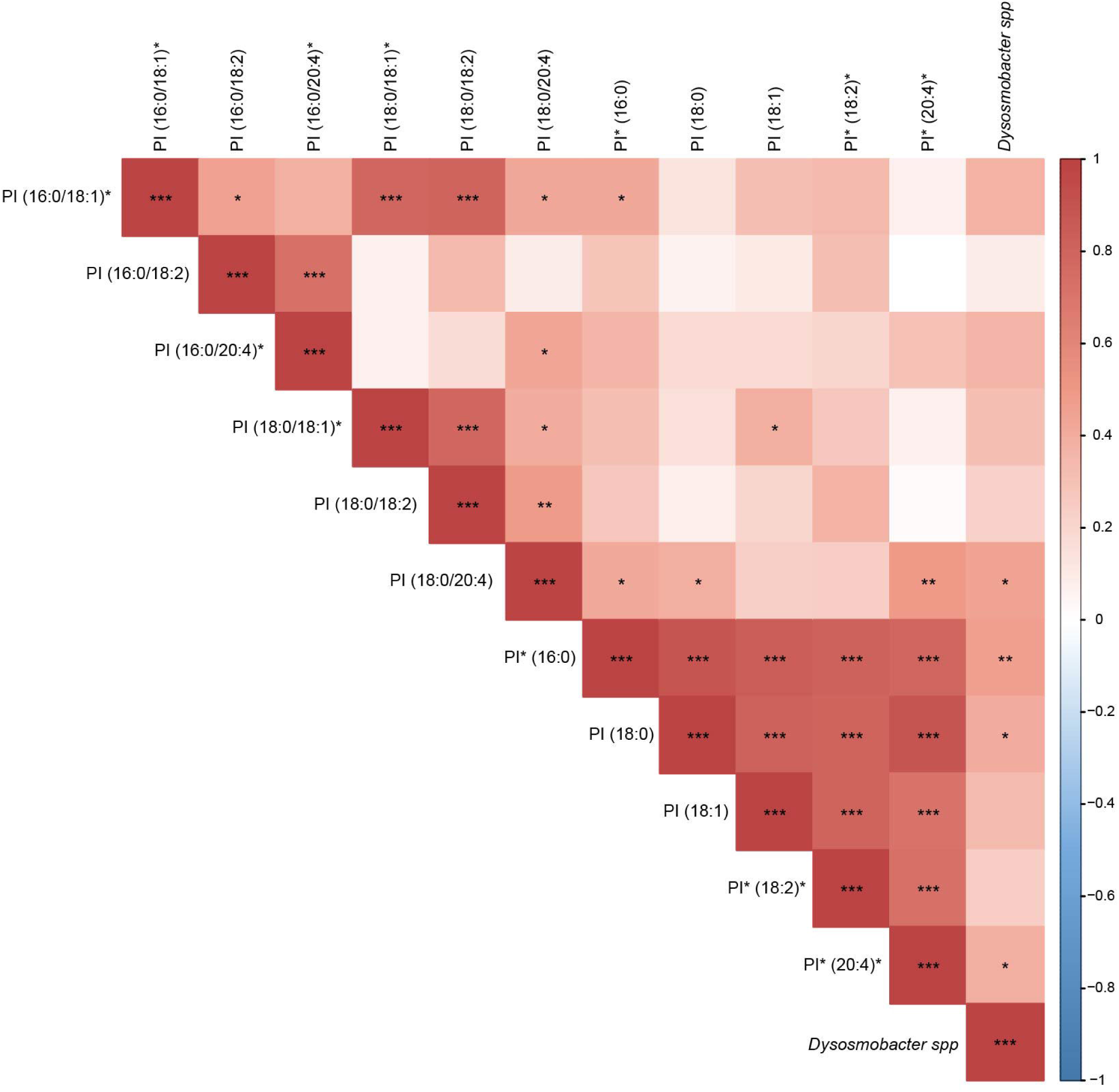
Spearman correlations between fecal Dysosmobacter spp, phosphatidylinositols and Lysophosphatidylinositols. Positive correlations are labeled in red and negative ones in blue. Significant correlations after FDR correction are labeled as follow, *: adjusted-p <0.05, **: adjusted-p <0.01, ***: adjusted-p <0.001. For clarity on the correlogram, PI and LysoPI names were shortened to only display the acyl chains composition.

PI accounts for ∼1% of total plasma lipids in humans (96) but their relationship with metabolic disorders is largely understudied. In 2019, a study in overweight and obese children reported reduced PI levels, suggesting a potential association with metabolic health (97). However, further research is needed to clarify these relationships.

### Lysoplasmalogens, phospholipids associated molecules and their potential role in metabolic health

In our study, fecal *Dysosmobacter spp* concentration was positively associated with GPC (P-16:0) a lysoplasmalogen molecule. However, literature on the role of lysoplasmalogens in obesity, T2D and metabolic syndrome is scarce, making it dìicult to draw definitive conclusions or propose clear hypotheses about their metabolic impact.

Additionally, glycerophosphorylcholine (GPC) and glycerophosphorylethanolamine (GPE) were also positively correlated with *Dysosmobacter spp. (****Table 2 and* *Figure 8*)**. These molecules are degradation products of (lyso)phospholipids, with GPC serving as a major circulating precursor of choline (98). However, few information regarding GPC & GPE association with metabolic disorders has been published. In this study, GPC and GPE were negatively correlated with plasma GLP-1, while GPC (P-16:0) was positively associated with plasma ADAMTS13, a metalloprotease involved in platelet aggregation regulation (99) ***(Table 3)***.

**Figure 8:**
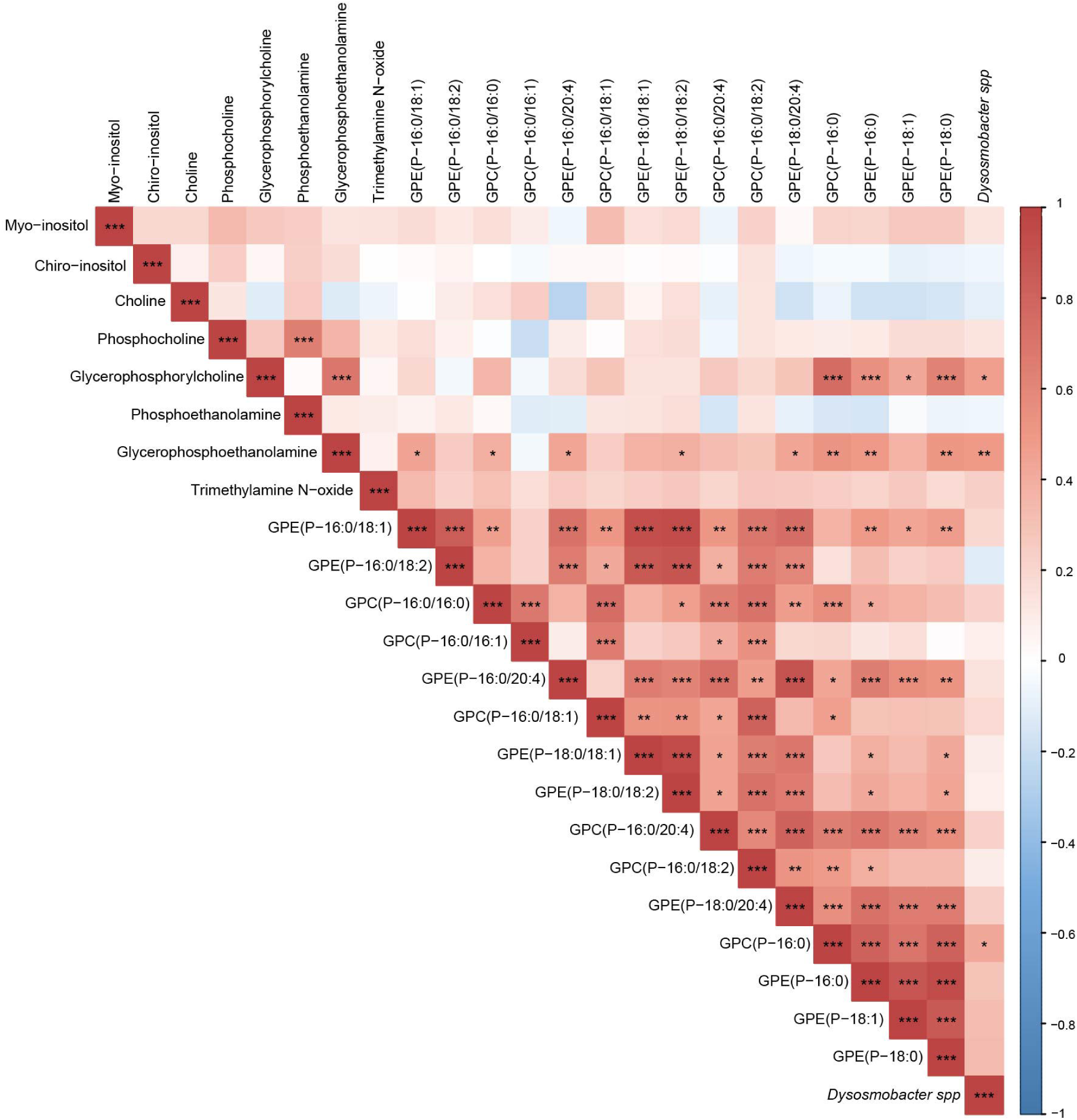
Spearman correlations between fecal Dysosmobacter spp, plasmalogens, lysoplasmalogens and phospholipid metabolism associated molecules. Positive correlations are labeled in red and negative ones in blue. Significant correlations after FDR correction are labeled as follow, *: adjusted-p <0.05, **: adjusted-p <0.01, ***: adjusted-p <0.001.

Although the limited available studies on the subject suggest that plasma PL, LysoPL and Lysoplasmalogens are altered in metabolic disorders, however, whether these changes are causative or consequential remains unclear and the mechanisms involved remain unknown. Variability between studies likely stems from differences in analytical methods, lipid subclass resolution, and cohort characteristics. Identifying consistent patterns in PL metabolism across studies is particularly challenging due to several factors. First, PLs represent a highly diverse family of molecules, and no single analytical method can comprehensively capture their full spectrum. Second, the sensitivity and resolution of lipidomic methods vary across research groups, limiting the ability to precisely characterize the acyl chain composition of detected phospholipids. These methodological inconsistencies complicate direct comparisons between studies and hinder a deeper understanding of PL dynamics in metabolic disorders.

### Association between Dysosmobacter spp and ceramides

A ceramide species (d18:1/20:0, d16:1/22:0, d20:1/18:0)* was positively correlated with *Dysosmobacter spp* fecal concentration (r-score = 0.512, adjusted p-value = 0.002) ***(Table 2 and* *Figure 9*)**. Ceramides are sphingolipids composed of a sphingoid base linked to a fatty acid via an amide bond and serve as precursors for more complex sphingolipids such as sphingomyelins and hexosylceramides. In plasma, ceramides are primarily transported by LDL and VLDL, suggesting a predominant hepatic origin, with a minor contribution from dietary sources (100).

**Figure 9:**
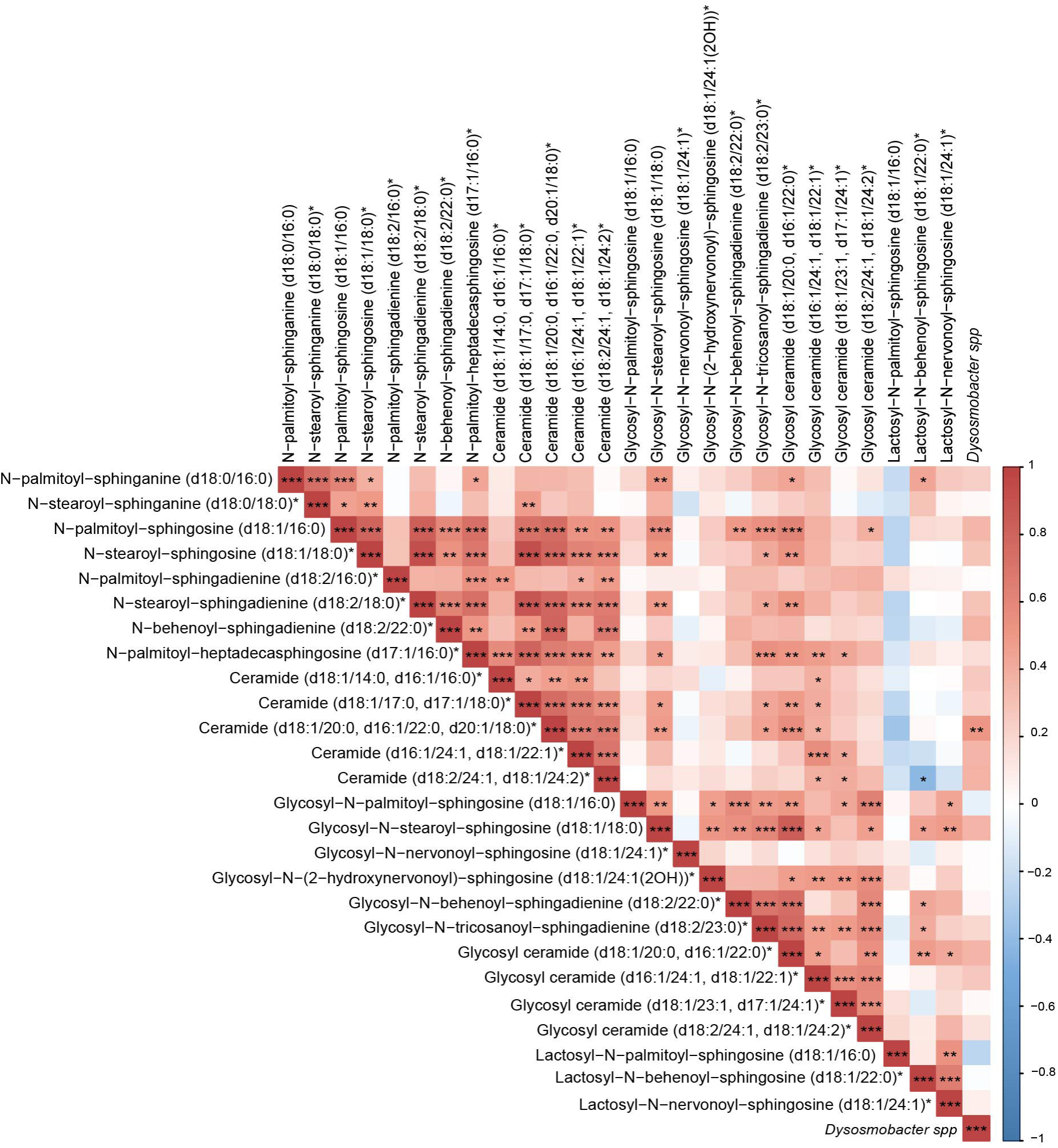
Spearman correlations between fecal Dysosmobacter spp and ceramides. Positive correlations are labeled in red and negative ones in blue. Significant correlations after FDR correction are labeled as follow, *: adjusted-p <0.05, **: adjusted-p <0.01, ***: adjusted-p <0.001.

In most studies ceramides are classified using the notation Cer(C:n) where “C” refers to the number of carbon in the fatty acid chain and “n” denotes the number of unsaturation. However, this convention often overlooks potential variations in the sphingoid base, which may influence the biological effects of these molecules.

As a lipid class, Ceramides (Cer), have been implicated in insulin resistance in animal models, though their role in human metabolism is still debated (29). They are frequently described as lipotoxic bioactive lipids that accumulate in the plasma of obese and insulin resistant individuals (101–105). In various cohorts across different countries, plasma levels of Cer16:0, Cer18:0, Cer20:0 and Cer22:0 have been linked to insulin resistance, HOMA-IR, β-cell function (HOMA-%S), incident diabetes and Matsuda index, and pro-inflammatory cytokines in individuals with CVD, as well as increased CVD incidence (82, 106–113). Unexpectedly, a 2018 study by Razquin et al. reported an inverse correlation between ceramides and T2D in two dìerent cohorts (87, 114). This contradictory finding may be explained by differences in lipid transport, with a sub-phenotype of patients exhibiting triglyceride-loaded LDL particles instead of ceramide-enriched LDL (87).

In our study, the analytical method did not provide sufficient resolution to differentiate between the three ceramide species grouped under the name *Ceramide (d18 :1/20 :0, d16 :1/22 :0, d20 :1/18 :0).* However, these species likely correspond to Cer20:0, Cer22:0 and Cer18:0, which have previously been associated with T2D, obesity and CVD markers. While we cannot determine the exact ceramide species linked to *Dysosmobacter spp*, all three potential molecules have been reported in association with adverse metabolic phenotypes, warranting further investigation into their role in host metabolism.

### Plasmatic cholesterol levels correlate with *Dysosmobacter spp*

Plasma total cholesterol levels positively correlated with *Dysosmobacter spp.* fecal concentrations, (r-score = 0.450; adjusted p-value = 0.014) ***(Table 2 and* *Figure 10*)**. While elevated total cholesterol is not a defining characteristic of metabolic syndrome, it is associated with an increased risk of CVD (115), particularly when levels exceed 200mg/dl. In this cohort, total cholesterol ranged from 102 to 312mg/dl, with some participants exceeding this threshold. The link between cholesterol and *Dysosmobacter welbionis*, is supported by a recent 2024 study by Li et al., which identified this bacterium as a cholesterol-metabolizing species. How this metabolic activity translates to plasma cholesterol levels, however, remains unclear (116).

**Figure 10:**
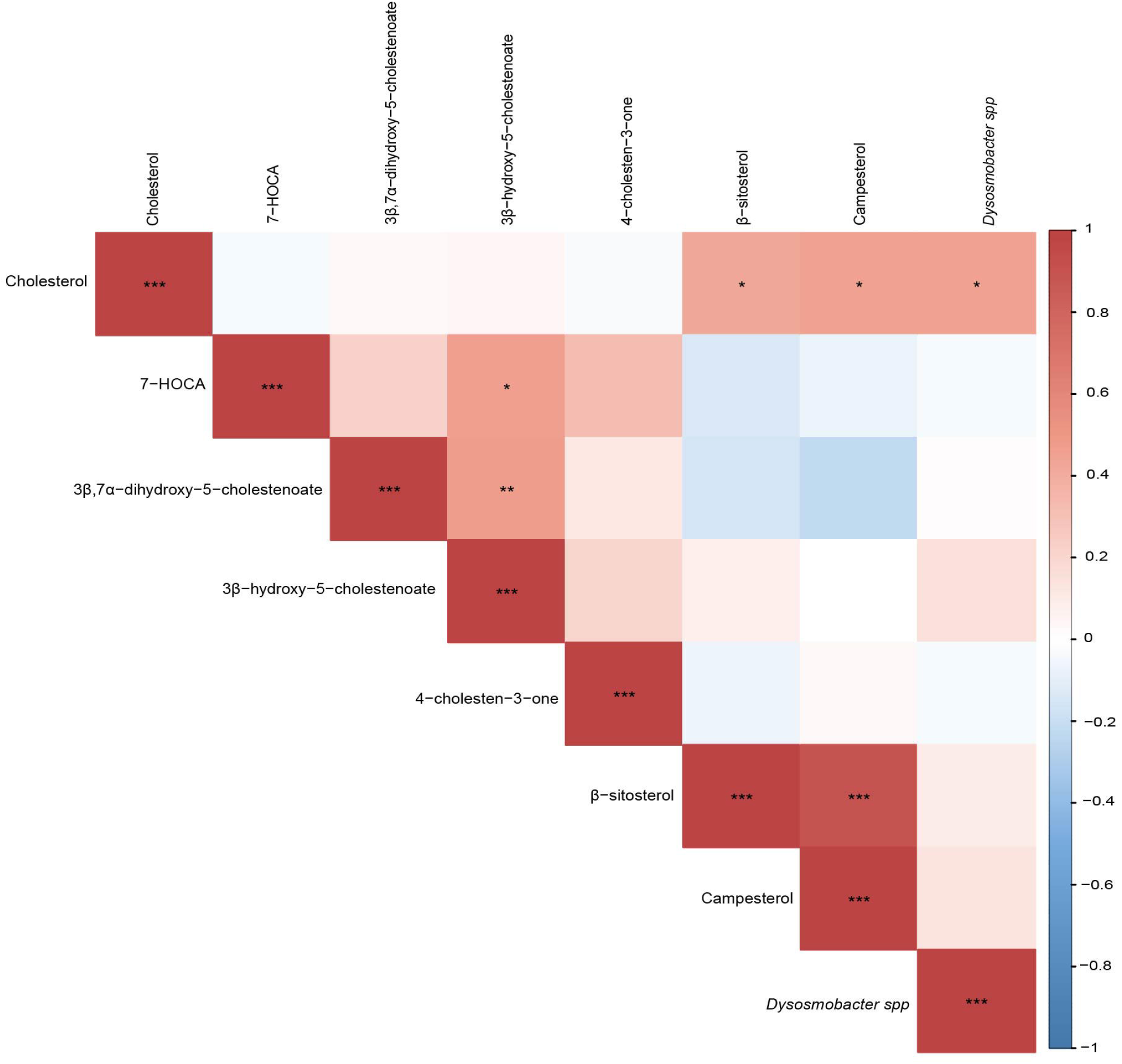
Spearman correlations between fecal Dysosmobacter spp and sterols. Positive correlations are labeled in red and negative ones in blue. Significant correlations after FDR correction are labeled as follow, *: adjusted-p <0.05, **: adjusted-p <0.01, ***: adjusted-p <0.001.

## Discussion

Interestingly, among the 862 identified metabolites covering multiple molecular families, only those belonging to the lipid super-pathway correlated significantly with *Dysosmobacter spp* fecal concentration. This finding aligns with a previous mouse study, where *Dysosmobacter welbionis* J115^T^ supplementation altered the lipid profiles in brown adipose tissue, colon and plasma under a high-fat diet (26). Specifically, six of nine detected acylcholines and eight of ten lysophosphatidylcholines (LysoPC) showed positive correlations with fecal *Dysosmobacter* levels, highlighting a potentially privileged relationship between this bacterium and host lipid metabolism. How *Dysosmobacter spp* modifies the lipid profile of various tissues, either directly or indirectly, remains unknow and requires further investigations.

Although *Dysosmobacter spp* itself was not directly associate with GLP-1 levels in this study, twelve lipid metabolites positively associated with *Dysosmobacter spp* fecal concentration were negatively correlated with GLP-1 plasma levels. GLP-1 is an incretin secreted by enteroendocrine cells in the intestine that plays a central role in glucose and energy homeostasis (28). Reduced circulating levels of this satiety-inducing hormone have been reported in obese and T2D patients, although findings are not consistent across all studies (117). Our previous research demonstrated that *D. welbionis* J115^T^ directly stimulated the dose-dependent secretion of GLP-1 in entero-endocrine cell *in vitro* (7). However, subsequent *in vivo* studies did not find elevated portal GLP-1 levels after *Dysosmobacter welbionis* J115^T^ supplementation, suggesting the involvement of complex mechanisms and potential indirect pathways linking this bacterium with host metabolism.

Overall, the results obtained in this human cohort support further exploration of *Dysosmobacter spp* and its potential influence on host lipid pathways, clarifying whether observed correlations represent causal mechanisms or indirect associations.

### Limitations of this study

Although our findings highlight previously undescribed links between Dysosmobacter spp. and the host lipidome, several limitations must be considered. Firstly, the study remains descriptive because participants were not specifically supplemented with *Dysosmobacter spp* to assess its effects. The human cohort was relatively small, included one timepoint and lacked a lean control group, restricting definitive causal interpretations.

Secondly, the metabolomics analysis provided only relative quantification of metabolites, without absolute reference values or a control group, limiting our ability to determine whether metabolite levels fall within physiological or pathological ranges and preventing direct comparisons to healthy populations.

Thirdly, due to methodological limitations, we could not clearly resolve all lipid species at the molecular level (e.g., certain ceramide species were grouped), hindering precise comparisons with existing literature.

Moreover, we cannot determine whether the observed lipids originated directly from host metabolism or from microbial metabolism. Gut bacteria can synthesize, accumulate and metabolize a wide variety of lipids, some identical to mammalian lipids and others unique, such as odd-chain fatty acids typically absent in those of mammal origin (118–120). Current knowledge regarding bacterial lipid composition remains limited, largely restricted to studies of model organisms such as *Escherichia coli*, and does not encompass the full diversity of the gut microbiota (121). Although certain bacterial lipids, including phospholipids with ethanolamine, choline, or inositol head groups, have been documented in various gut microbiota species (119, 121–124), there is very little known about the microbial production of MAGs, DAGs, ceramides, or acylcholines (122, 124–126). How much of these lipids interact with the host is still unclear, as we don’t know which subclasses can be absorbed by the host to integrate its own metabolism. Notably, recent evidence demonstrates that odd-chain bacterial lipids, including sphingolipids produced by *Bacteroides thetaiotaomicron*, can be detected in the gut epithelium and portal vein, although not in the liver, suggesting active absorption and subsequent metabolism by host tissues (127). While in the referenced study the lipids were administered to the mice via gavage as purified solution rather than produced *in situ* by bacteria, it remains plausible that similar microbial lipids could be naturally synthesized and absorbed within the gut. Additionally, gut bacteria might also influence the host lipid profile indirectly through modulation of host metabolic pathways (121, 128–135). Therefore, *Dysosmobacter spp*. could influence host metabolism either directly, through microbial lipid production and absorption, or indirectly, by shaping overall microbial community dynamics that subsequently impact host lipid profiles. These limitations highlight the necessity of future studies that investigate both the direct lipidomic profiles of gut bacteria and the causal relationships between specific bacterial taxa, host lipid metabolism, and metabolic health outcomes.

## Conclusion

*Dysosmobacter welbionis* is a recently discovered gut bacterium strongly linked to beneficial metabolic effects, positioning it as an attractive “next-generation” probiotic candidate for addressing obesity and diabetes. To effectively translate these promising findings, however, it remains crucial to understand how this bacterium interacts with host metabolism; the present study advances this goal by providing novel and valuable insights by demonstrating that previously unexplored microbial-host interaction and offering a foundation for exploring potential underlying mechanisms.

## Declarations

### Ethics approval and consent to participate

Institutional Review Board Statement: The study was conducted according to the guidelines of the Declaration of Helsinki, and approved by the Ethics Committee Commission d’Ethique Biomédicale Hospitalo-facultaire of the Université catholique de Louvain (Brussel, Belgium). The study was registered at https://clinicaltrials.gov as trial no. NCT02637115 (protocol code: 2015/02JUL/369, approved on the 13 July 2015). Informed consent was obtained from all subjects involved in the study.

### Consent for publication

Not applicable.

### Availabilities of data and materials

The datasets used and/or analyzed during the current study are available from the corresponding author on reasonable request.

### Authors’ contribution

Conceptualization: CP, MVH, PDC; Data curation: CP, CD; Formal analysis: CP; Funding acquisition: PDC; Investigation: CP, CD, AE; Methodology: CP, CD, AE, MVH, PDC; Project administration: MVH, PDC; Resources: PDC and NMD; Supervision: MVH, PDC; Validation: MVH, PDC; Vizualisation: CP; Writing original draft: CP; Writing – review & editing: CP, AE, MVH, PDC; all authors agreed with the final submitted manuscript.

## Acknowledgments

We thank the team involved in the initial collection of fecal and blood samples: de Barsy M, Druart C., Loumaye A., Maiter D., Thissen J-P and Hermans MP.

## List of abbreviations

20-HETE: 20-hydroxyeicosatetraenoic acid
2-palmitoleoyl-GPC* (16:1)*: PC* (16:1)* [2]: 1-palmitoleoyl-GPC* (16:1)* = PC* (16:1)* [1]
AChE: Acetylcholinesterase
AlkP: Alkaline phosphatase
ALT: Alanine transaminase
AST: Aspartate transaminase
BChE: Butyrylcholinesterase
BMI: Body Mass Index
BW: Bodyweight
CB1: Cannabinoid receptor-1
Cer: Ceramide
CK: Creatinine kinase
COX: Cyclooxygenase
CVD: Cardiovascular diseases
CYP450: Cytochrome P450
DAG (16:0/18:1) [2]*: Palmitoyl-oleoyl-glycerol (16:0/18:1) [2]*
DAG (16:0/18:2) [2]*: Palmitoyl-linoleoyl-glycerol (16:0/18:2) [2]*
DAG (16:0/20:4) [1]*: Palmitoyl-arachidonoyl-glycerol (16:0/20:4) [1]*
DAG (16:1/18:1) [1]*: Palmitoleoyl-oleoyl-glycerol (16:1/18:1) [1]*
DAG (16:1/18:2) [1]*: Palmitoleoyl-linoleoyl-glycerol (16:1/18:2) [1]*
DAG (18:0/20:4) [1]*: Stearoyl-arachidonoyl-glycerol (18:0/20:4) [1]*
DAG (18:0/20:4) [2]*: Stearoyl-arachidonoyl-glycerol (18:0/20:4) [2]*
DAG (18:1/18:2) [1]: Oleoyl-linoleoyl-glycerol (18:1/18:2) [1]
DAG (18:1/18:2) [2]: Oleoyl-linoleoyl-glycerol (18:1/18:2) [1]
DAG (18:1/18:3) [2]*: Oleoyl-linolenoyl-glycerol (18:1/18:3) [2]*
DAG (18:1/20:4) [1]*: Oleoyl-arachidonoyl-glycerol (18:1/20:4) [1]*
DAG (18:1/20:4) [2]*: Oleoyl-arachidonoyl-glycerol (18:1/20:4) [2]*
DAG (18:2/18:2) [1]*: Linoleoyl-linoleoyl-glycerol (18:2/18:2) [1]*
DAG (18:2/18:2) [2]*: Linoleoyl-linoleoyl-glycerol (18:2/18:2) [2]*
DAG (18:2/18:3) [1]*: Linoleoyl-linolenoyl-glycerol (18:2/18:3) [1]*
DAG (18:2/18:3) [2]*: Linoleoyl-linolenoyl-glycerol (18:2/18:3) [2]*
DAG (18:2/20:4) [1]*: Linoleoyl-arachidonoyl-glycerol (18:2/20:4) [1]*
DAG (18:2/20:4) [2]*: Linoleoyl-arachidonoyl-glycerol (18:2/20:4) [2]*
DAG (18:2/22:6) [2]*: Linoleoyl-docosahexaenoyl-glycerol (18:2/22:6) [2]*
DAG(14:0/18:2) [1]*: Myristoyl-linoleoyl-glycerol (14:0/18:2) [1]*
DAG(16:0/18:1) [1]*: Palmitoyl-oleoyl-glycerol (16:0/18:1) [1]*
DAG(16:0/18:2) [1]*: Palmitoyl-linoleoyl-glycerol (16:0/18:2) [1]*
DAG(16:0/18:3) [2]*: Palmitoyl-linolenoyl-glycerol (16:0/18:3) [2]*
DAG(16:0/20:4) [2]*: Palmitoyl-arachidonoyl-glycerol (16:0/20:4) [2]*
DAG(18:1/18:1) [2]*: Oleoyl-oleoyl-glycerol (18:1/18:1) [2]*
DAG(18:1/18:1) [1]*: Oleoyl-oleoyl-glycerol (18:1/18:1) [1]*
DAG: Diacylglycerol
DDP-4: Dipeptidyl peptidase 4
EET: Epoxyeicosatrienoic acids
FDR: False discovery rate
GDF-15: Growth differentiation factor 15
GFR: Glomerular filtration rate
GGT: Gamma-glutamyl transferase
GIP: Gastric inhibitory polypeptide
GLP-1: Glucagon-like-peptide-1
GPC: Glycerophosphorylcholine
GPE: Glycerophosphorylethanolamine
HDL: High density lipoprotein
HFD: High-fat diet
hsCRP: C-reactive protein
IFABP: Intestinal fatty-acid binding protein
IL-6: Interleukin-6
IP-10: Interferon gamma-induced protein 10
IR: Insulin resistance
LBP: Lipopolysaccharide binding protein
LDH: Lactate dehydrogenase
LOX: Lipoxygenase
LPS: Lipopolycaccharide
LXA4: Lipoxin A4
LysoPC: Lysophosphocholine
LysoPE: Lysophosphatidylethanolamine
LysoPI: Lysophosphatidylinositol
LysoPL: Lysophospholipid
MAG: Monoacylglycerol
MCP-1: Plasmatic monocyte chemoattractant protein-1
MDC: Macrophage derived chemokine
nAchR: Nicotinic acetylcholine receptor
NEFA: Plasma non-esterified fatty acids
ND: Normal diet
PC (14:0/16:0): 1-myristoyl-2-palmitoyl-GPC (14:0/16:0)
PC (14:0/20:4)*: 1-myristoyl-2-arachidonoyl-GPC (14:0/20:4)*
PC (16:0/16:0): 1,2-dipalmitoyl-GPC (16:0/16:0)
PC (16:0/16:1)*: 1-palmitoyl-2-palmitoleoyl-GPC (16:0/16:1)*
PC (16:0/18:0): 1-palmitoyl-2-stearoyl-GPC (16:0/18:0)
PC (16:0/18:1): 1-palmitoyl-2-oleoyl-GPC (16:0/18:1)
PC (16:0/18:2): 1-palmitoyl-2-linoleoyl-GPC (16:0/18:2)
PC (16:0/18:3n3)*: 1-palmitoyl-2-alpha-linolenoyl-GPC (16:0/18:3n3)*
PC (16:0/18:3n6)*: 1-palmitoyl-2-gamma-linolenoyl-GPC (16:0/18:3n6)*
PC (16:0/20:3n3 or 6)*: 1-palmitoyl-2-dihomo-linolenoyl-GPC (16:0/20:3n3 or 6)*
PC (16:0/20:4n6): 1-palmitoyl-2-arachidonoyl-GPC (16:0/20:4n6)
PC (16:0/22:6): 1-palmitoyl-2-docosahexaenoyl-GPC (16:0/22:6)
PC (16:1/18:2)*: 1-palmitoleoyl-2-linoleoyl-GPC (16:1/18:2)*
PC (16:1/18:3)*: 1-palmitoleoyl-2-linolenoyl-GPC (16:1/18:3)*
PC (18:0/18:0): 1,2-distearoyl-GPC (18:0/18:0)
PC (18:0/18:1): 1-stearoyl-2-oleoyl-GPC (18:0/18:1)
PC (18:0/18:2)*: 1-stearoyl-2-linoleoyl-GPC (18:0/18:2)*
PC (18:0/20:4): 1-stearoyl-2-arachidonoyl-GPC (18:0/20:4)
PC (18:0/22:6): 1-stearoyl-2-docosahexaenoyl-GPC (18:0/22:6)
PC (18:1): 1-oleoyl-GPC (18:1)
PC (18:1/18:2)*: 1-oleoyl-2-linoleoyl-GPC (18:1/18:2)*
PC (18:1/22:6)*: 1-oleoyl-2-docosahexaenoyl-GPC (18:1/22:6)*
PC (18:2): 1-linoleoyl-GPC (18:2)
PC (18:2/18:2): 1,2-dilinoleoyl-GPC (18:2/18:2)
PC (18:2/18:3)*: 1-linoleoyl-2-linolenoyl-GPC (18:2/18:3)*
PC (18:2/20:4n6)*: 1-linoleoyl-2-arachidonoyl-GPC (18:2/20:4n6)*
PC (18:3)*: 1-linoleoyl-GPC (18:3)*
PC (24:0): 1-lignoceroyl-GPC (24:0)
PC* (16:0)*: 1-palmitoyl-GPC (16:0) = PC (16:0), 2-palmitoyl-GPC* (16:0)*
PC* (20:4)*: 1-arachidonoyl-GPC* (20:4)*
PC: Phosphocholine
PE (16:0): 1-palmitoyl-GPE (16:0)
PE (16:0/18:1): 1-palmitoyl-2-oleoyl-GPE (16:0/18:1)
PE (16:0/18:2): 1-palmitoyl-2-linoleoyl-GPE (16:0/18:2)
PE (16:0/20:4)*: 1-palmitoyl-2-arachidonoyl-GPE (16:0/20:4)*
PE (16:0/22:6)*: 1-palmitoyl-2-docosahexaenoyl-GPE (16:0/22:6)*
PE (18:0) [1]: 1-stearoyl-GPE (18:0)
PE (18:0)*: 2-stearoyl-GPE (18:0)* [2]
PE (18:0/18:1): 1-stearoyl-2-oleoyl-GPE (18:0/18:1)
PE (18:0/18:2)*: 1-stearoyl-2-linoleoyl-GPE (18:0/18:2)*
PE (18:0/20:4): 1-stearoyl-2-arachidonoyl-GPE (18:0/20:4)
PE (18:0/22:6)*: 1-stearoyl-2-docosahexaenoyl-GPE (18:0/22:6)*
PE (18:1): 1-oleoyl-GPE (18:1)
PE (18:1/18:2)*: 1-oleoyl-2-linoleoyl-GPE (18:1/18:2)*
PE (18:1/20:4)*: 1-oleoyl-2-arachidonoyl-GPE (18:1/20:4)*
PE (18:1/22:6)*: 1-oleoyl-2-docosahexaenoyl-GPE (18:1/22:6)*
PE (18:2)*: 1-linoleoyl-GPE (18:2)*
PE (18:2/18:2)*: 1,2-dilinoleoyl-GPE (18:2/18:2)*
PE: Phosphatidylethanolamine
PI (16:0/18:1)*: 1-palmitoyl-2-oleoyl-GPI (16:0/18:1)*
PI (16:0/18:2): 1-palmitoyl-2-linoleoyl-GPI (16:0/18:2)
PI (16:0/20:4): 1-palmitoyl-2-arachidonoyl-GPI (16:0/20:4)*
PI (18:0): 1-stearoyl-GPI (18:0)
PI (18:0/18:1)*: 1-stearoyl-2-oleoyl-GPI (18:0/18:1)*
PI (18:0/18:2): 1-stearoyl-2-linoleoyl-GPI (18:0/18:2)
PI (18:0/20:4): 1-stearoyl-2-arachidonoyl-GPI (18:0/20:4)
PI (18:1): 1-oleoyl-GPI (18:1)
PI* (16:0): 1-palmitoyl-GPI* (16:0)
PI* (18:2)*: 1-linoleoyl-GPI* (18:2)*
PI* (20:4)*: 1-arachidonoyl-GPI* (20:4)*
PI: Phosphatidylinositol
PL: Phospholipid
pNA: p-nitroanilide
PUFA: Polyunsaturated fatty acid
PYY: Peptide YY
sCD40L: Soluble CD40 ligand
siCAM-1: Soluble intercellular adhesion molecule-1
svCAM-1: Soluble vascular cell adhesion molecule-1
T2D: Type 2 diabetes
TG: Triglyceride
WBC: White blood cell count

## References

1. Rubino F, Cummings DE, Eckel RH, Cohen RV, Wilding JPH, Brown WA, et al. Definition and diagnostic criteria of clinical obesity. Lancet Diabetes Endocrinol. 2025;13(3):221–62.

2. Kassi E, Pervanidou P, Kaltsas G, Chrousos G. Metabolic syndrome: definitions and controversies. BMC Med. 2011;9:48.

3. Gilijamse PW, Hartstra AV, Levin E, Wortelboer K, Serlie MJ, Ackermans MT, et al. Treatment with Anaerobutyricum soehngenii: a pilot study of safety and dose-response effects on glucose metabolism in human subjects with metabolic syndrome. NPJ Biofilms Microbiomes. 2020;6(1):16.

4. Depommier C, Everard A, Druart C, Plovier H, Van Hul M, Vieira-Silva S, et al. Supplementation with Akkermansia muciniphila in overweight and obese human volunteers: a proof-of-concept exploratory study. Nat Med. 2019;25(7):1096–103.

5. Le Roy T, Moens de Hase E, Van Hul M, Paquot A, Pelicaen R, Régnier M, et al. Dysosmobacter welbionis is a newly isolated human commensal bacterium preventing diet-induced obesity and metabolic disorders in mice. Gut. 2022;71(3):534–43.

6. Le Roy T, Van der Smissen P, Paquot A, Delzenne N, Muccioli GG, Collet JF, et al. Dysosmobacter welbionis gen. nov., sp. nov., isolated from human faeces and emended description of the genus Oscillibacter. Int J Syst Evol Microbiol. 2020;70(9):4851–8.

7. Moens de Hase E, Neyrinck AM, Rodriguez J, Cnop M, Paquot N, Thissen J-P, et al. Impact of metformin and Dysosmobacter welbionis on diet-induced obesity and diabetes: from clinical observation to preclinical intervention. Diabetologia. 2024;67(2):333–45.

8. Aderemi AV, Ayeleso AO, Oyedapo OO, Mukwevho E. Metabolomics: A Scoping Review of Its Role as a Tool for Disease Biomarker Discovery in Selected Non-Communicable Diseases. Metabolites. 2021;11(7).

9. Johnson CH, Ivanisevic J, Siuzdak G. Metabolomics: beyond biomarkers and towards mechanisms. Nature Reviews Molecular Cell Biology. 2016;17(7):451–9.

10. Payab M, Tayanloo-Beik A, Falahzadeh K, Mousavi M, Salehi S, Djalalinia S, et al. Metabolomics prospect of obesity and metabolic syndrome; a systematic review. J Diabetes Metab Disord. 2022;21(1):889–917.

11. Rangel-Huerta OD, Pastor-Villaescusa B, Gil A. Are we close to defining a metabolomic signature of human obesity? A systematic review of metabolomics studies. Metabolomics. 2019;15(6):93.

12. Visconti A, Le Roy CI, Rosa F, Rossi N, Martin TC, Mohney RP, et al. Interplay between the human gut microbiome and host metabolism. Nat Commun. 2019;10(1):4505.

13. Wilmanski T, Rappaport N, Earls JC, Magis AT, Manor O, Lovejoy J, et al. Blood metabolome predicts gut microbiome alpha-diversity in humans. Nat Biotechnol. 2019;37(10):1217–28.

14. Asnicar F, Berry SE, Valdes AM, Nguyen LH, Piccinno G, Drew DA, et al. Microbiome connections with host metabolism and habitual diet from 1,098 deeply phenotyped individuals. Nat Med. 2021;27(2):321–32.

15. Partula V, Deschasaux-Tanguy M, Mondot S, Victor-Bala A, Bouchemal N, Lécuyer L, et al. Associations between untargeted plasma metabolomic signatures and gut microbiota composition in the Milieu Intérieur population of healthy adults. British Journal of Nutrition. 2021;126(7):982–92.

16. Dekkers KF, Sayols-Baixeras S, Baldanzi G, Nowak C, Hammar U, Nguyen D, et al. An online atlas of human plasma metabolite signatures of gut microbiome composition. Nat Commun. 2022;13(1):5370.

17. Zhang Y, Chen R, Zhang D, Qi S, Liu Y. Metabolite interactions between host and microbiota during health and disease: Which feeds the other? Biomed Pharmacother. 2023;160:114295.

18. Depommier C, Flamand N, Pelicaen R, Maiter D, Thissen JP, Loumaye A, et al. Linking the Endocannabinoidome with Specific Metabolic Parameters in an Overweight and Insulin-Resistant Population: From Multivariate Exploratory Analysis to Univariate Analysis and Construction of Predictive Models. Cells. 2021;10(1).

19. Depommier C, Everard A, Druart C, Maiter D, Thissen JP, Loumaye A, et al. Serum metabolite profiling yields insights into health promoting effect of A. muciniphila in human volunteers with a metabolic syndrome. Gut Microbes. 2021;13(1):1994270.

20. Wallace TM, Levy JC, Matthews DR. Use and abuse of HOMA modeling. Diabetes Care. 2004;27(6):1487–95.

21. Dennis JM, Shields BM, Hill AV, Knight BA, McDonald TJ, Rodgers LR, et al. Precision Medicine in Type 2 Diabetes: Clinical Markers of Insulin Resistance Are Associated With Altered Short- and Long-term Glycemic Response to DPP-4 Inhibitor Therapy. Diabetes Care. 2018;41(4):705–12.

22. Babu H, Sperk M, Ambikan AT, Rachel G, Viswanathan VK, Tripathy SP, et al. Plasma Metabolic Signature and Abnormalities in HIV-Infected Individuals on Long-Term Successful Antiretroviral Therapy. Metabolites. 2019;9(10).

23. Depommier C, Vitale RM, Iannotti FA, Silvestri C, Flamand N, Druart C, et al. Beneficial Effects of Akkermansia muciniphila Are Not Associated with Major Changes in the Circulating Endocannabinoidome but Linked to Higher Mono-Palmitoyl-Glycerol Levels as New PPARalpha Agonists. Cells. 2021;10(1).

24. William R. Procedures for psychological, psychometric, and personality research. Package ‘pscyh; 2021.

25. Wei T, Simko V, Levy M, Xie Y, Jin Y, Zemla J, et al. corrplot: Visualization of a Correlation Matrix (0.92). WWW document] URL https://github.com/taiyun/corrplot [accessed 23 September …; 2021.

26. Moens De Hase E, Petitfils C, Alhouayek M, Depommier C, Le Faouder P, Delzenne NM, et al. Dysosmobacter welbionis effects on glucose, lipid, and energy metabolism are associated with specific bioactive lipids. Journal of Lipid Research. 2023;64(10):100437.

27. Henderson GC. Plasma Free Fatty Acid Concentration as a Modifiable Risk Factor for Metabolic Disease. Nutrients. 2021;13(8):2590.

28. Drucker DJ. Mechanisms of Action and Therapeutic Application of Glucagon-like Peptide-1. Cell Metab. 2018;27(4):740–56.

29. Tonks KT, Coster AC, Christopher MJ, Chaudhuri R, Xu A, Gagnon-Bartsch J, et al. Skeletal muscle and plasma lipidomic signatures of insulin resistance and overweight/obesity in humans. Obesity (Silver Spring, Md). 2016;24(4):908–16.

30. Kulkarni H, Mamtani M, Wong G, Weir JM, Barlow CK, Dyer TD, et al. Genetic correlation of the plasma lipidome with type 2 diabetes, prediabetes and insulin resistance in Mexican American families. BMC Genetics. 2017;18(1):48.

31. Poursharifi P, Madiraju SRM, Prentki M. Monoacylglycerol signalling and ABHD6 in health and disease. Diabetes Obes Metab. 2017;19 Suppl 1:76–89.

32. Vangipurapu J, Fernandes Silva L, Kuulasmaa T, Smith U, Laakso M. Microbiota-Related Metabolites and the Risk of Type 2 Diabetes. Diabetes Care. 2020;43(6):1319–25.

33. Yousri NA, Suhre K, Yassin E, Al-Shakaki A, Robay A, Elshafei M, et al. Metabolic and Metabo-Clinical Signatures of Type 2 Diabetes, Obesity, Retinopathy, and Dyslipidemia. Diabetes. 2022;71(2):184–205.

34. Muccioli GG, Naslain D, Bäckhed F, Reigstad CS, Lambert DM, Delzenne NM, et al. The endocannabinoid system links gut microbiota to adipogenesis. Molecular Systems Biology. 2010;6(1):392.

35. Mehrpouya-Bahrami P, Chitrala KN, Ganewatta MS, Tang C, Murphy EA, Enos RT, et al. Blockade of CB1 cannabinoid receptor alters gut microbiota and attenuates inflammation and diet-induced obesity. Sci Rep. 2017;7(1):15645.

36. Naja K, Anwardeen N, Al-Hariri M, Al Thani AA, Elrayess MA. Pharmacometabolomic Approach to Investigate the Response to Metformin in Patients with Type 2 Diabetes: A Cross-Sectional Study. Biomedicines. 2023;11(8):2164.

37. Audet-Delage Y, Villeneuve L, Grégoire J, Plante M, Guillemette C. Identification of Metabolomic Biomarkers for Endometrial Cancer and Its Recurrence after Surgery in Postmenopausal Women. Frontiers in Endocrinology. 2018;9:87.

38. Zeleznik OA, Poole EM, Lindstrom S, Kraft P, Van Hylckama Vlieg A, Lasky-Su JA, et al. Metabolomic analysis of 92 pulmonary embolism patients from a nested case-control study identifies metabolites associated with adverse clinical outcomes. Journal of thrombosis and haemostasis: JTH. 2018;16(3):500–7.

39. Vorkas PA, Shalhoub J, Isaac G, Want EJ, Nicholson JK, Holmes E, et al. Metabolic Phenotyping of Atherosclerotic Plaques Reveals Latent Associations between Free Cholesterol and Ceramide Metabolism in Atherogenesis. Journal of Proteome Research. 2015;14(3):1389–99.

40. Heresi GA, Mey JT, Bartholomew JR, Haddadin IS, Tonelli AR, Dweik RA, et al. Plasma metabolomic profile in chronic thromboembolic pulmonary hypertension. Pulmonary Circulation. 2020;10(1):2045894019890553.

41. Germain A, Barupal DK, Levine SM, Hanson MR. Comprehensive Circulatory Metabolomics in ME/CFS Reveals Disrupted Metabolism of Acyl Lipids and Steroids. Metabolites. 2020;10(1):34.

42. Kang JW, Tang X, Walton CJ, Brown MJ, Brewer RA, Maddela RL, et al. Multi-Omic Analyses Reveal Bifidogenic Effect and Metabolomic Shifts in Healthy Human Cohort Supplemented With a Prebiotic Dietary Fiber Blend. Frontiers in Nutrition. 2022;9:908534.

43. Zarei I, Oppel RC, Borresen EC, Brown RJ, Ryan EP. Modulation of plasma and urine metabolome in colorectal cancer survivors consuming rice bran. Integrative Food, Nutrition and Metabolism. 2019;6(3).

44. Li C, Bundy JD, Tian L, Zhang R, Chen J, Kelly TN, et al. Examination of serum metabolome altered by dietary carbohydrate, milk protein, and soy protein interventions identified novel metabolites associated with blood pressure: the ProBP trial. Molecular nutrition & food research. 2023;67(20):e2300044.

45. Zambrana LE, Weber AM, Borresen EC, Zarei I, Perez J, Perez C, et al. Daily Rice Bran Consumption for 6 Months Influences Serum Glucagon-Like Peptide 2 and Metabolite Profiles without Differences in Trace Elements and Heavy Metals in Weaning Nicaraguan Infants at 12 Months of Age. Current Developments in Nutrition. 2021;5(9):nzab101.

46. Hunt R, de Mortemer Taveau R. The effects of a number of derivatives of choline and analogous compounds on the blood-pressure: US Government Printing Office; 1911.

47. Akimov MG, Kudryavtsev DS, Kryukova EV, Fomina-Ageeva EV, Zakharov SS, Gretskaya NM, et al. Arachidonoylcholine and Other Unsaturated Long-Chain Acylcholines Are Endogenous Modulators of the Acetylcholine Signaling System. Biomolecules. 2020;10(2):283.

48. Jain R, Kutty KM, Huang SN, Kean K. Pseudocholinesterase/high-density lipoprotein cholesterol ratio in serum of normal persons and of hyperlipoproteinemics. Clinical Chemistry. 1983;29(6):1031–3.

49. Iwasaki T, Yoneda M, Nakajima A, Terauchi Y. Serum Butyrylcholinesterase is Strongly Associated with Adiposity, the Serum Lipid Profile and Insulin Resistance. Internal Medicine. 2007;46(19):1633–9.

50. Randell EW, Mathews MS, Zhang H, Seraj JS, Sun G. Relationship between serum butyrylcholinesterase and the metabolic syndrome. Clin Biochem. 2005;38(9):799–805.

51. De Vriese C, Gregoire F, Lema-Kisoka R, Waelbroeck M, Robberecht P, Delporte C. Ghrelin degradation by serum and tissue homogenates: identification of the cleavage sites. Endocrinology. 2004;145(11):4997–5005.

52. Lafontan M, Langin D. Lipolysis and lipid mobilization in human adipose tissue. Progress in Lipid Research. 2009;48(5):275–97.

53. Zhang Y, Liu Y, Sun J, Zhang W, Guo Z, Ma Q. Arachidonic acid metabolism in health and disease. MedComm. 2023;4(5):e363.

54. Sonnweber T, Pizzini A, Nairz M, Weiss G, Tancevski I. Arachidonic Acid Metabolites in Cardiovascular and Metabolic Diseases. International Journal of Molecular Sciences. 2018;19(11):3285.

55. Blood exams referential Liège CHU - Urea [Available from: https://www.chu.ulg.ac.be/jcms/c_498724/fr/uree-sang.

56. Galgani JE, Aguirre CA, Uauy RD, Díaz EO. Plasma Arachidonic Acid Influences Insulin-Stimulated Glucose Uptake in Healthy Adult Women. Annals of Nutrition and Metabolism. 2007;51(5):482–9.

57. Mustonen A-M, Nieminen P. Dihomo-γ-Linolenic Acid (20:3n-6)—Metabolism, Derivatives, and Potential Significance in Chronic Inflammation. International Journal of Molecular Sciences. 2023;24(3):2116.

58. Das UN. “Cell Membrane Theory of Senescence” and the Role of Bioactive Lipids in Aging, and Aging Associated Diseases and Their Therapeutic Implications. Biomolecules. 2021;11(2).

59. Borgeson E, Johnson AM, Lee YS, Till A, Syed GH, Ali-Shah ST, et al. Lipoxin A4 Attenuates Obesity-Induced Adipose Inflammation and Associated Liver and Kidney Disease. Cell Metab. 2015;22(1):125–37.

60. Yang J, Wang M, Yang D, Yan H, Wang Z, Yan D, et al. Integrated lipids biomarker of the prediabetes and type 2 diabetes mellitus Chinese patients. Frontiers in Endocrinology. 2023;13.

61. Rauschert S, Uhl O, Koletzko B, Kirchberg F, Mori TA, Huang R-C, et al. Lipidomics Reveals Associations of Phospholipids With Obesity and Insulin Resistance in Young Adults. The Journal of Clinical Endocrinology and Metabolism. 2016;101(3):871–9.

62. Pang S-J, Liu T-T, Pan J-C, Man Q-Q, Song S, Zhang J. The Association between the Plasma Phospholipid Profile and Insulin Resistance: A Population-Based Cross-Section Study from the China Adult Chronic Disease and Nutrition Surveillance. Nutrients. 2024;16(8):1205.

63. Sigruener A, Kleber ME, Heimerl S, Liebisch G, Schmitz G, Maerz W. Glycerophospholipid and sphingolipid species and mortality: the Ludwigshafen Risk and Cardiovascular Health (LURIC) study. PLoS One. 2014;9(1):e85724.

64. Alshehry ZH, Mundra PA, Barlow CK, Mellett NA, Wong G, McConville MJ, et al. Plasma Lipidomic Profiles Improve on Traditional Risk Factors for the Prediction of Cardiovascular Events in Type 2 Diabetes Mellitus. Circulation. 2016;134(21):1637–50.

65. Birgbauer E, Chun J. New developments in the biological functions of lysophospholipids. Cell Mol Life Sci. 2006;63(23):2695–701.

66. Ishii I, Fukushima N, Ye X, Chun J. Lysophospholipid receptors: signaling and biology. Annu Rev Biochem. 2004;73:321–54.

67. D’Arrigo P, Servi S. Synthesis of lysophospholipids. Molecules. 2010;15(3):1354–77.

68. Ha CY, Kim JY, Paik JK, Kim OY, Paik Y-H, Lee EJ, et al. The association of specific metabolites of lipid metabolism with markers of oxidative stress, inflammation and arterial stiffness in men with newly diagnosed type 2 diabetes. Clinical Endocrinology. 2012;76(5):674–82.

69. Kuang A, Erlund I, Herder C, Westerhuis JA, Tuomilehto J, Cornelis MC. Lipidomic Response to Coffee Consumption. Nutrients. 2018;10(12):1851.

70. Dommel S, Bluher M. Does C-C Motif Chemokine Ligand 2 (CCL2) Link Obesity to a Pro-Inflammatory State? Int J Mol Sci. 2021;22(3).

71. Rull A, Camps J, Alonso-Villaverde C, Joven J. Insulin resistance, inflammation, and obesity: role of monocyte chemoattractant protein-1 (or CCL2) in the regulation of metabolism. Mediators Inflamm. 2010;2010.

72. Qiao Q, Bouwman FG, van Baak MA, Roumans NJT, Vink RG, Mariman ECM. Plasma Levels of Triglycerides and IL-6 Are Associated With Weight Regain and Fat Mass Expansion. J Clin Endocrinol Metab. 2022;107(7):1920–9.

73. Menezes AMB, Oliveira PD, Wehrmeister FC, Goncalves H, Assuncao MCF, Tovo-Rodrigues L, et al. Association between interleukin-6, C-reactive protein and adiponectin with adiposity: Findings from the 1993 pelotas (Brazil) birth cohort at 18 and 22 years. Cytokine. 2018;110:44–51.

74. Vozarova B, Weyer C, Hanson K, Tataranni PA, Bogardus C, Pratley RE. Circulating interleukin-6 in relation to adiposity, insulin action, and insulin secretion. Obes Res. 2001;9(7):414–7.

75. Bastard JP, Jardel C, Bruckert E, Blondy P, Capeau J, Laville M, et al. Elevated levels of interleukin 6 are reduced in serum and subcutaneous adipose tissue of obese women after weight loss. J Clin Endocrinol Metab. 2000;85(9):3338–42.

76. Webber M, Krishnan A, Thomas NG, Cheung BM. Association between serum alkaline phosphatase and C-reactive protein in the United States National Health and Nutrition Examination Survey 2005-2006. Clin Chem Lab Med. 2010;48(2):167–73.

77. Kim JH, Lee HS, Park HM, Lee YJ. Serum alkaline phosphatase level is positively associated with metabolic syndrome: A nationwide population-based study. Clin Chim Acta. 2020;500:189–94.

78. Sigvardsen CM, Richter MM, Engelbeen S, Kleinert M, Richter EA. GDF15 is still a mystery hormone. Trends Endocrinol Metab. 2024.

79. Huynh K, Barlow CK, Jayawardana KS, Weir JM, Mellett NA, Cinel M, et al. High-Throughput Plasma Lipidomics: Detailed Mapping of the Associations with Cardiometabolic Risk Factors. Cell Chemical Biology. 2019;26(1):71–84.e4.

80. Mika A, Sledzinski T. Alterations of specific lipid groups in serum of obese humans: a review. Obes Rev. 2017;18(2):247–72.

81. del Bas JM, Caimari A, Rodriguez-Naranjo MI, Childs CE, Paras Chavez C, West AL, et al. Impairment of lysophospholipid metabolism in obesity: altered plasma profile and desensitization to the modulatory properties of n–3 polyunsaturated fatty acids in a randomized controlled trial12. The American Journal of Clinical Nutrition. 2016;104(2):266–79.

82. Weir JM, Wong G, Barlow CK, Greeve MA, Kowalczyk A, Almasy L, et al. Plasma lipid profiling in a large population-based cohort. Journal of Lipid Research. 2013;54(10):2898–908.

83. Pikó P, Pál L, Szűcs S, Kósa Z, Sándor J, Ádány R. Obesity-Related Changes in Human Plasma Lipidome Determined by the Lipidyzer Platform. Biomolecules. 2021;11(2):326.

84. Bellot P, Moia MN, Reis BZ, Pedrosa LFC, Tasic L, Barbosa F, Jr., et al. Are Phosphatidylcholine and Lysophosphatidylcholine Body Levels Potentially Reliable Biomarkers in Obesity? A Review of Human Studies. Mol Nutr Food Res. 2023;67(7):e2200568.

85. Barber MN, Risis S, Yang C, Meikle PJ, Staples M, Febbraio MA, et al. Plasma lysophosphatidylcholine levels are reduced in obesity and type 2 diabetes. PloS One. 2012;7(7):e41456.

86. Park S, Sadanala KC, Kim E-K. A Metabolomic Approach to Understanding the Metabolic Link between Obesity and Diabetes. Molecules and Cells. 2015;38(7):587–96.

87. Razquin C, Toledo E, Clish CB, Ruiz-Canela M, Dennis C, Corella D, et al. Plasma Lipidomic Profiling and Risk of Type 2 Diabetes in the PREDIMED Trial. Diabetes Care. 2018;41(12):2617–24.

88. Wang-Sattler R, Yu Z, Herder C, Messias AC, Floegel A, He Y, et al. Novel biomarkers for pre-diabetes identified by metabolomics. Molecular Systems Biology. 2012;8(1):615.

89. Ferrannini E, Natali A, Camastra S, Nannipieri M, Mari A, Adam K-P, et al. Early metabolic markers of the development of dysglycemia and type 2 diabetes and their physiological significance. Diabetes. 2013;62(5):1730–7.

90. Suvitaival T, Bondia-Pons I, Yetukuri L, Pöhö P, Nolan JJ, Hyötyläinen T, et al. Lipidome as a predictive tool in progression to type 2 diabetes in Finnish men. Metabolism. 2018;78:1–12.

91. Soga T, Ohishi T, Matsui T, Saito T, Matsumoto M, Takasaki J, et al. Lysophosphatidylcholine enhances glucose-dependent insulin secretion via an orphan G-protein-coupled receptor. Biochem Biophys Res Commun. 2005;326(4):744–51.

92. Orešič M, Hyötyläinen T, Kotronen A, Gopalacharyulu P, Nygren H, Arola J, et al. Prediction of non-alcoholic fatty-liver disease and liver fat content by serum molecular lipids. Diabetologia. 2013;56(10):2266–74.

93. Kühn T, Floegel A, Sookthai D, Johnson T, Rolle-Kampczyk U, Otto W, et al. Higher plasma levels of lysophosphatidylcholine 18:0 are related to a lower risk of common cancers in a prospective metabolomics study. BMC Medicine. 2016;14(1):13.

94. García-Fontana B, Morales-Santana S, Díaz Navarro C, Rozas-Moreno P, Genilloud O, Vicente Pérez F, et al. Metabolomic profile related to cardiovascular disease in patients with type 2 diabetes mellitus: A pilot study. Talanta. 2016;148:135–43.

95. Fiehn O, Garvey WT, Newman JW, Lok KH, Hoppel CL, Adams SH. Plasma metabolomic profiles reflective of glucose homeostasis in non-diabetic and type 2 diabetic obese African-American women. PLoS One. 2010;5(12):e15234.

96. Xia T, Ren H, Zhang W, Xia Y. Lipidome-Wide Characterization of Phosphatidylinositols and Phosphatidylglycerols on C=C Location Level. Analytica chimica acta. 2020;1128:107–15.

97. Anjos S, Feiteira E, Cerveira F, Melo T, Reboredo A, Colombo S, et al. Lipidomics Reveals Similar Changes in Serum Phospholipid Signatures of Overweight and Obese Pediatric Subjects. Journal of Proteome Research. 2019;18(8):3174–83.

98. Aisyah R, Ohshima N, Watanabe D, Nakagawa Y, Sakuma T, Nitschke F, et al. GDE5/Gpcpd1 activity determines phosphatidylcholine composition in skeletal muscle and regulates contractile force in mice. Commun Biol. 2024;7(1):604.

99. Shaw RJ, Abrams ST, Badu S, Toh CH, Dutt T. The Highs and Lows of ADAMTS13 Activity. J Clin Med. 2024;13(17).

100. Larsen PJ, Tennagels N. On ceramides, other sphingolipids and impaired glucose homeostasis. Molecular Metabolism. 2014;3(3):252–60.

101. Meikle PJ, Wong G, Barlow CK, Weir JM, Greeve MA, MacIntosh GL, et al. Plasma Lipid Profiling Shows Similar Associations with Prediabetes and Type 2 Diabetes. PLOS ONE. 2013;8(9):e74341.

102. Summers SA. Ceramides in insulin resistance and lipotoxicity. Progress in Lipid Research. 2006;45(1):42–72.

103. Huang H, Kasumov T, Gatmaitan P, Heneghan HM, Kashyap SR, Schauer PR, et al. Gastric bypass surgery reduces plasma ceramide subspecies and improves insulin sensitivity in severely obese patients. Obesity (Silver Spring). 2011;19(11):2235–40.

104. Samad F, Hester KD, Yang G, Hannun YA, Bielawski J. Altered adipose and plasma sphingolipid metabolism in obesity: a potential mechanism for cardiovascular and metabolic risk. Diabetes. 2006;55(9):2579–87.

105. Hanamatsu H, Ohnishi S, Sakai S, Yuyama K, Mitsutake S, Takeda H, et al. Altered levels of serum sphingomyelin and ceramide containing distinct acyl chains in young obese adults. Nutr Diabetes. 2014;4(10):e141.

106. Boon J, Hoy AJ, Stark R, Brown RD, Meex RC, Henstridge DC, et al. Ceramides contained in LDL are elevated in type 2 diabetes and promote inflammation and skeletal muscle insulin resistance. Diabetes. 2013;62(2):401–10.

107. de Mello VD, Lankinen M, Schwab U, Kolehmainen M, Lehto S, Seppanen-Laakso T, et al. Link between plasma ceramides, inflammation and insulin resistance: association with serum IL-6 concentration in patients with coronary heart disease. Diabetologia. 2009;52(12):2612–5.

108. Ichi I, Nakahara K, Miyashita Y, Hidaka A, Kutsukake S, Inoue K, et al. Association of ceramides in human plasma with risk factors of atherosclerosis. Lipids. 2006;41(9):859–63.

109. Holland WL, Summers SA. Sphingolipids, insulin resistance, and metabolic disease: new insights from in vivo manipulation of sphingolipid metabolism. Endocr Rev. 2008;29(4):381–402.

110. Kopprasch S, Dheban S, Schuhmann K, Xu A, Schulte KM, Simeonovic CJ, et al. Detection of Independent Associations of Plasma Lipidomic Parameters with Insulin Sensitivity Indices Using Data Mining Methodology. PLoS One. 2016;11(10):e0164173.

111. Bae JC, Wander PL, Lemaitre RN, Fretts AM, Sitlani CM, Bui HH, et al. Associations of plasma sphingolipids with measures of insulin sensitivity, β-cell function, and incident diabetes in Japanese Americans. Nutrition, Metabolism and Cardiovascular Diseases. 2024;34(3):633–41.

112. Haus JM, Kashyap SR, Kasumov T, Zhang R, Kelly KR, Defronzo RA, et al. Plasma ceramides are elevated in obese subjects with type 2 diabetes and correlate with the severity of insulin resistance. Diabetes. 2009;58(2):337–43.

113. Chen L, Dong Y, Bhagatwala J, Raed A, Huang Y, Zhu H. Vitamin D3 Supplementation Increases Long-Chain Ceramide Levels in Overweight/Obese African Americans: A Post-Hoc Analysis of a Randomized Controlled Trial. Nutrients. 2020;12(4):981.

114. Razquin C, Liang L, Toledo E, Clish CB, Ruiz-Canela M, Zheng Y, et al. Plasma lipidome patterns associated with cardiovascular risk in the PREDIMED trial: A case-cohort study. Int J Cardiol. 2018;253:126–32.

115. Jung E, Kong SY, Ro YS, Ryu HH, Shin SD. Serum Cholesterol Levels and Risk of Cardiovascular Death: A Systematic Review and a Dose-Response Meta-Analysis of Prospective Cohort Studies. Int J Environ Res Public Health. 2022;19(14).

116. Li C, Stražar M, Mohamed AMT, Pacheco JA, Walker RL, Lebar T, et al. Gut microbiome and metabolome profiling in Framingham heart study reveals cholesterol-metabolizing bacteria. Cell. 2024;187(8):1834–52.e19.

117. Holst JJ. Annual Prize Lecture 2024: Endogenous physiological mechanisms as basis for the treatment of obesity and type 2 diabetes. J Physiol. 2024;602(24):6613–29.

118. Ryan E, Joyce SA, Clarke DJ. Membrane lipids from gut microbiome-associated bacteria as structural and signalling molecules. Microbiology. 2023;169(3).

119. Lamichhane S, Sen P, Alves MA, Ribeiro HC, Raunioniemi P, Hyötyläinen T, et al. Linking Gut Microbiome and Lipid Metabolism: Moving beyond Associations. Metabolites. 2021;11(1):55.

120. Rezanka T, Sigler K. Odd-numbered very-long-chain fatty acids from the microbial, animal and plant kingdoms. Prog Lipid Res. 2009;48(3-4):206–38.

121. Brown EM, Clardy J, Xavier RJ. Gut microbiome lipid metabolism and its impact on host physiology. Cell Host & Microbe. 2023;31(2):173–86.

122. López-Lara IM, Geiger O. Bacterial lipid diversity. Biochimica et Biophysica Acta (BBA) - Molecular and Cell Biology of Lipids. 2017;1862(11):1287–99.

123. Lee T-H, Charchar P, Separovic F, E. Reid G, Yarovsky I, Aguilar M-I. The intricate link between membrane lipid structure and composition and membrane structural properties in bacterial membranes. Chemical Science. 2024;15(10):3408–27.

124. Geiger O, Sohlenkamp C, López-Lara IM. Formation of Bacterial Glycerol-Based Membrane Lipids: Pathways, Enzymes, and Reactions. In: Geiger O, editor. Biogenesis of Fatty Acids, Lipids and Membranes. Cham: Springer International Publishing; 2019. p. 87–107.

125. Ganesh BP, Hall A, Ayyaswamy S, Nelson JW, Fultz R, Major A, et al. Diacylglycerol kinase synthesized by commensal Lactobacillus reuteri diminishes protein kinase C phosphorylation and histamine-mediated signaling in the mammalian intestinal epithelium. Mucosal Immunology. 2018;11(2):380–93.

126. Vulevic J, McCartney AL, Gee JM, Johnson IT, Gibson GR. Microbial species involved in production of 1,2-sn-diacylglycerol and effects of phosphatidylcholine on human fecal microbiota. Appl Environ Microbiol. 2004;70(9):5659–66.

127. Johnson EL, Heaver SL, Waters JL, Kim BI, Bretin A, Goodman AL, et al. Sphingolipids produced by gut bacteria enter host metabolic pathways impacting ceramide levels. Nature Communications. 2020;11(1):2471.

128. Manca C, Boubertakh B, Leblanc N, Deschênes T, Lacroix S, Martin C, et al. Germ-free mice exhibit profound gut microbiota-dependent alterations of intestinal endocannabinoidome signaling. Journal of Lipid Research. 2020;61(1):70–85.

129. Schoeler M, Caesar R. Dietary lipids, gut microbiota and lipid metabolism. Reviews in Endocrine and Metabolic Disorders. 2019;20(4):461–72.

130. Fu J, Bonder MJ, Cenit MC, Tigchelaar EF, Maatman A, Dekens JAM, et al. The Gut Microbiome Contributes to a Substantial Proportion of the Variation in Blood Lipids. Circulation Research. 2015;117(9):817–24.

131. Druart C, Bindels LB, Schmaltz R, Neyrinck AM, Cani PD, Walter J, et al. Ability of the gut microbiota to produce PUFA-derived bacterial metabolites: Proof of concept in germ-free versus conventionalized mice. Molecular Nutrition & Food Research. 2015;59(8):1603–13.

132. Kishino S, Takeuchi M, Park S-B, Hirata A, Kitamura N, Kunisawa J, et al. Polyunsaturated fatty acid saturation by gut lactic acid bacteria àecting host lipid composition. Proceedings of the National Academy of Sciences. 2013;110(44):17808–13.

133. Kindt A, Liebisch G, Clavel T, Haller D, Hörmannsperger G, Yoon H, et al. The gut microbiota promotes hepatic fatty acid desaturation and elongation in mice. Nature Communications. 2018;9(1):3760.

134. Velagapudi VR, Hezaveh R, Reigstad CS, Gopalacharyulu P, Yetukuri L, Islam S, et al. The gut microbiota modulates host energy and lipid metabolism in mice. Journal of Lipid Research. 2010;51(5):1101–12.

135. Albouery M, Buteau B, Grégoire S, Cherbuy C, Pais de Barros J-P, Martine L, et al. Age-Related Changes in the Gut Microbiota Modify Brain Lipid Composition. Frontiers in Cellular and Infection Microbiology. 2020;9:444.

